# HR3/RORα-mediated cholesterol sensing regulates TOR signaling

**DOI:** 10.1101/2024.04.09.588676

**Authors:** Mette Lassen, Keith Pardee, Lisa H. Pedersen, Olga Kubrak, Takashi Koyama, Sasha Necakov, Suya Liu, Arnis Kuksis, Gilles Lajoie, Aled Edwards, Aurelio A. Teleman, Martin R. Larsen, Henry M. Krause, Michael J. Texada, Kim Rewitz

## Abstract

Cells and organisms adjust their growth based on the availability of cholesterol, which is essential for cellular functions. However, the mechanisms by which cells sense cholesterol levels and translate these into growth signals are not fully understood. We report that cholesterol rapidly activates the master growth-regulatory TOR pathway in *Drosophila* tissues. We identify the nuclear receptor HR3, an ortholog of mammalian RORα, as an essential factor in cholesterol-induced TOR activation. We demonstrate that HR3 binds cholesterol and promotes TOR pathway activation through a non-genomic mechanism acting upstream of the Rag GTPases. Similarly, we find that RORα is necessary for cholesterol-mediated TOR activation in human cells, suggesting that HR3/RORα represents a conserved mechanism for cholesterol sensing that couples cholesterol availability to TOR-pathway activity. These findings advance our understanding of how cholesterol influences cell growth, with implications for cholesterol-related diseases and cancer.

**Highlights:**

- Cholesterol leads to dynamic TOR pathway activation, driving systemic growth
- HR3 in *Drosophila* binds cholesterol and couples its availability to TOR activation
- HR3 acts upstream of Rag GTPases to activate TOR in response to lysosomal cholesterol
- Mammalian HR3 ortholog RORα is required for cholesterol-induced TOR activation

## Introduction

Cholesterol is widely recognized as a fundamental component of cellular membranes. Beyond its structural role as a lipid, it functions as a critical signaling molecule, affecting signaling pathways and physiological processes. Cholesterol can interact with proteins through specific cholesterol-binding domains, thereby influencing various cellular processes such as signal transduction^1–4^. Furthermore, cholesterol is the precursor to other signaling molecules, including steroid hormones, bile acids, and oxysterols, that can activate specific receptors and modulate gene expression^5,6^. Dysregulated cholesterol levels have been implicated in many health disorders ranging from cardiovascular diseases to promoting cancerous growth^7,8^. The deep involvement of cholesterol in physiology highlights the importance of cellular mechanisms that sense cholesterol availability and in response modulate cellular growth.

In mammalian cells, which can synthesize sterols *de novo*, cholesterol homeostasis is governed by sterol-sensing mechanisms within the endoplasmic reticulum (ER). Through regulation of the transcription factor Sterol Response Element Binding Protein (SREBP), cholesterol biosynthesis and cellular uptake are modulated to maintain lipid homeostasis^9^. The demand for cholesterol increases during rapid cell proliferation – for example, during development and in cancer cells. To meet this demand, upregulation of pathways such as phosphatidylinositol 3-kinase (PI3K)/Protein Kinase B (PKB/AKT) enhances the uptake of exogenous cholesterol^10^. Although cholesterol is necessary for cell-membrane synthesis, increased cholesterol availability also directly drives cell growth and fuels rapid proliferation in cancers. Unlike the well-characterized SREBP pathway, the cellular mechanisms by which cholesterol abundance or deficiency is linked to growth-regulatory pathways are poorly defined. We recently demonstrated that cholesterol stimulates systemic growth during development through the activation of insulin signaling in *Drosophila*^11^. While our genetic evidence implicated the target of rapamycin (TOR) pathway in this process, the cholesterol-induced TOR responses and the molecular mechanisms linking cholesterol to TOR activation – particularly identifying the components that directly interact with cholesterol to register its availability – remained elusive. Thus, the mechanisms by which cellular cholesterol levels are conveyed to TOR for translation into growth signals are poorly understood. The TOR pathway integrates environmental cues including nutrients, oxygen availability, growth stimuli, and cellular stressors into a single signal that regulates cellular growth^12,13^. Activated TOR promotes cellular growth and protein synthesis by promoting ribosomal biogenesis and enhancing translation while simultaneously inhibiting autophagy and regulating metabolism. In mammalian cell culture, cholesterol has emerged as a key factor that activates the TOR pathway^3,14^. Extracellular cholesterol is taken up through endocytosis and trafficked to the late endosome/lysosome compartment. When cholesterol accumulates within these organelles, the TOR complex is recruited by a large protein assembly to the lysosomal membrane, where it interacts with the small GTPase Rheb, which in turn activates TOR kinase activity. In conditions of nutrient scarcity, including low cholesterol, TOR remains localized to the cytosol. In this compartment, TOR has little access to Rheb and therefore cannot be activated.

A large network of proteins orchestrates the nutrient-responsive localization of TOR to the lysosome. The lysosomal recruitment of TOR is ultimately governed by small GTPases of the Rag family^12,15^. These proteins associate with the LAMTOR1 subunit of the Ragulator complex, which anchors them to the lysosomal membrane. In response to nutrient abundance, a diverse array of mechanisms configures the GTP/GDP loading of Rag heterodimers in such way that they bind to RAPTOR, a component of the TOR complex, which maintains TOR in association with the lysosomal surface, where it can be activated by Rheb. Rag heterodimers are active and can interact with RAPTOR when RagA/B is guanosine triphosphate (GTP)-bound and RagC/D is guanosine diphosphate (GDP)- bound^12^. This balance is maintained by GTPase-activating protein (GAP) complexes such as GATOR1 and guanine nucleotide exchange factor (GEF) complexes including Ragulator. In nutrient-depleted states, GATOR1 induces GTP hydrolysis on RagA, leading to loss of TOR activation. This mechanism is influenced by amino acids, which affect the GAP activity of GATOR1. Low amino-acid levels also negatively regulate Ragulator, preventing it from activating the Rag GTPases through its GEF functionality.

In contrast to the level of detail with which amino acid-mediated TOR activation is understood, the cholesterol-sensing mechanisms that are capable of directly binding to or detecting intracellular cholesterol and interacting with the TOR pathway are poorly defined. Multiple intracellular cholesterol pools are present, including those in lysosomes, deposited across the plasma membrane and ER, as well as at membrane contact sites^16^. Given the variability in cholesterol concentration across these distinct cellular compartments, it is likely that several distinct cholesterol-sensing mechanisms exist, each specifically designed to detect different cholesterol pools and independently modulate the TOR pathway. Recent reports suggest that a mechanism by which lysosomal cholesterol activates the TOR complex in mammalian cells involves LYCHOS, a G protein–coupled receptor (GPCR)-like protein integral to the lysosomal membrane that modulates the Rag nucleotide-binding state in a cholesterol-dependent manner via interaction with GATOR1^14^. Additionally, the mammalian lysosomal transmembrane protein SLC38A9 is important for the activation of TOR in response to both cholesterol and amino acids^3^. In *Drosophila*, dietary cholesterol activates TOR, with increasing levels driving rapid growth and development^11^. Intracellular cholesterol accumulation in lysosomal compartments caused by deficiency in the cholesterol transporter Niemann-Pick Type C1 (NPC1) drives strong activation of the TOR pathway in both *Drosophila* and mammals^3,11^. This common feature suggests that the mechanisms coupling cholesterol levels to the activation of the TOR pathway are evolutionarily ancient and conserved. *Drosophila* possesses an ortholog of LYCHOS, named Anchor, but an SLC38A9-like protein does not appear to exist in flies, implying that other ancient and possibly conserved pathways exist that detect cholesterol and link its availability to the activation of TOR signaling.

Nuclear receptors bind lipophilic molecules and exert their regulatory influence through both direct effects on transcription (genomic effects) and more immediate signaling mechanisms that do not involve changes in gene transcription (non-genomic effects), orchestrating a diverse range of cellular processes^17,18^. DHR96, a *Drosophila* nuclear receptor involved in cholesterol homeostasis, binds cholesterol and regulates the expression of genes involved in cholesterol uptake, metabolism, and transport^1,4^. In mammals, the nuclear receptor Retinoic acid receptor (RAR)-related Orphan Receptor alpha (RORα) influences cholesterol homeostasis by modulating the transcription of genes important for cholesterol synthesis, uptake, and efflux^19^. However, whether nuclear receptors bind cholesterol and transduce levels of this lipid to modulate the TOR pathway is not known.

Here, we demonstrate that dietary cholesterol intake leads to rapid and dynamic activation of the TOR pathway in tissues of *Drosophila*. This response is modulated by the *Drosophila* RORα ortholog, HR3, which – like RORα – binds cholesterol and is activated by this ligand. Although HR3 is known to be transcriptionally upregulated by the ecdysone steroid hormone EcR, our results reveal that HR3 regulates growth through the TOR pathway in response to cholesterol independently of ecdysone-mediated effects. This regulation involves rapid cholesterol-induced TOR activation that in part is independent of the transcriptional functions of HR3, through an isoform of HR3 that lacks a DNA-binding domain (DBD). Reducing *HR3* levels in cells mitigate the overactivation of TOR caused by the intralysosomal accumulation of cholesterol resulting from depletion of *Npc1*a. This indicates that HR3 is necessary for TOR activation by lysosomal cholesterol. Our findings suggest that HR3 activates the TOR pathway upstream of Rag proteins. Furthermore, our findings indicate that RORα is involved in cholesterol-induced TOR activation in human cells, suggesting that a conserved function of HR3/RORα is to couple cholesterol levels to TOR-pathway activation.

## Results

### Cholesterol induces TOR-pathway activity

Dietary intake of cholesterol and its subsequent cellular absorption from the bloodstream are key for cellular cholesterol acquisition in both *Drosophila* and mammals. Cholesterol esters carried by lipoprotein particles are internalized from circulating lipoproteins through endocytosis and are transported via the endosomal pathway to lysosomes, where they are hydrolyzed so that free (unesterified) cholesterol can be trafficked between hydrophobic compartments of the cell^20,21^. Unlike mammals, which can also synthesize cholesterol *de novo*, *Drosophila* (and other insects) lack a functional sterol-biosynthesis pathway. Consequently, cholesterol availability in *Drosophila* can be precisely manipulated via dietary modifications ^22^. We used a synthetic diet^23^ that allows cholesterol supplementation at a range of doses, which promotes larval growth and development^11,24^. We specifically examined the role of dietary cholesterol during the last larval instar, a period characterized by rapid growth driven by nutrient intake. To investigate the dose-dependent effects on TOR-pathway activity, we supplemented the diet with cholesterol at concentrations ranging from 1.2 µg/mL to 80 µg/mL, reflecting ecologically relevant sterol levels^25,26^. Larvae were initially cholesterol-depleted by transferring them to synthetic food containing 0 µg/mL cholesterol for 10 hours at 84 hours after egg laying (AEL), then re-fed with synthetic food containing different doses of cholesterol for 6 hours (until 100 h AEL). Dietary cholesterol dose-dependently activated TOR signaling in fat-body tissue, as indicated by increased phosphorylation of Ribosomal protein S6 (pS6) and of its activator S6 kinase (pS6K), a direct target of TOR, reflecting TOR pathway activity^27^ (Figures 1A and 1B). Higher cholesterol doses correlated with elevated whole-body protein levels, suggesting that cholesterol promotes organismal protein synthesis and growth (Figure 1C), consistent with TOR activation.

**Figure 1:**
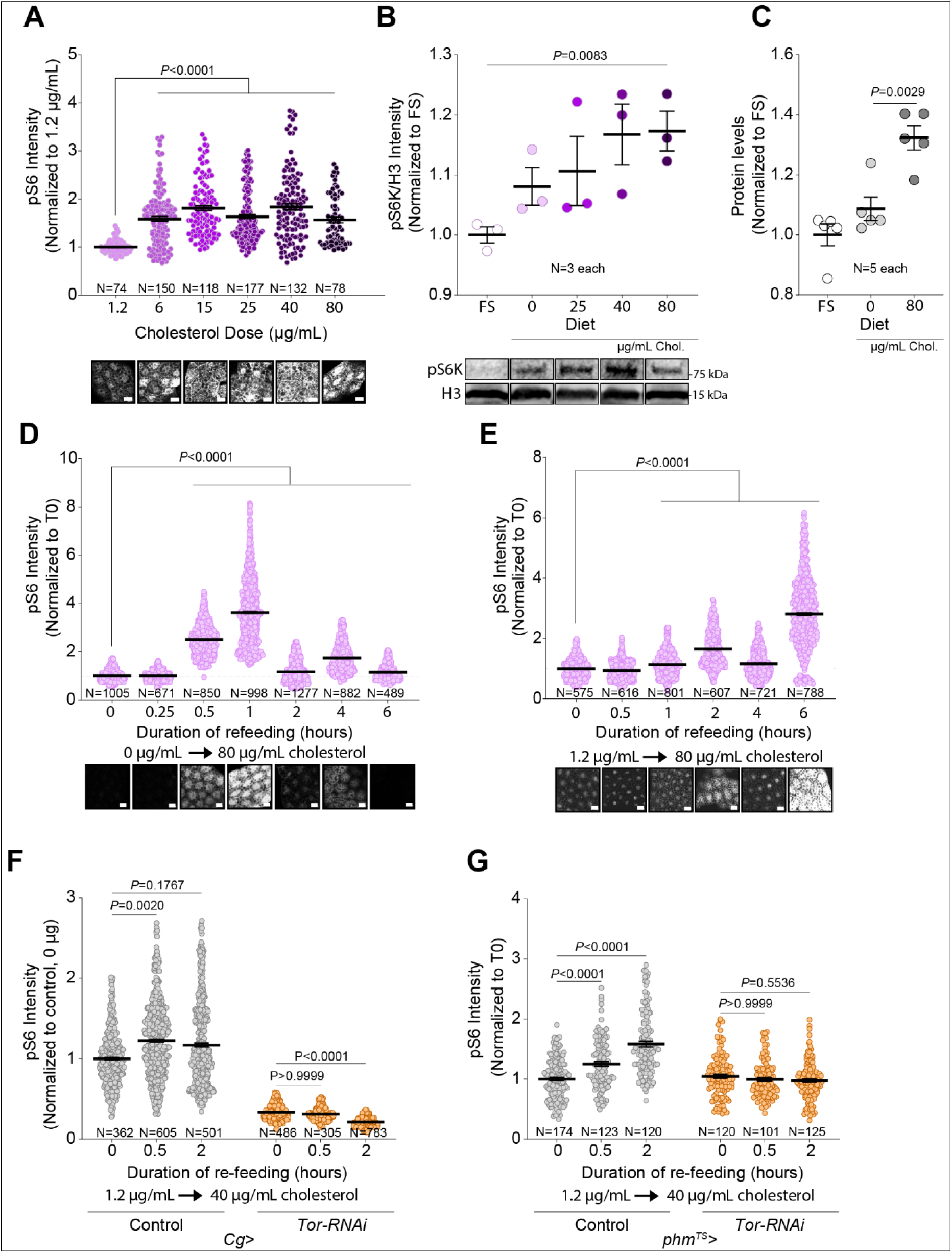
Cholesterol availability alters TOR activity in the fat-body and prothoracic gland of *Drosophila*. (A) Quantification of cholesterol-dose sensitivity of the TOR pathway using pS6 immunohistochemistry staining in larval fat body, with representative images (scale bar 20, µm). (B) Dose response, with representative blot images, of whole-larval pS6, normalized to histone H3, 1 hour after refeeding following 10 hours’ cholesterol deprivation. FS, full starvation. (C) Measurement of whole-larval protein levels by BCA after 4 hours’ refeeding following cholesterol deprivation. FS, full starvation; 0, cholesterol deprivation (0 µg/mL); 80, 80 µg/mL cholesterol. (D) Dynamics of fat-body pS6 staining levels (with representative images, 20-µm scale bar) during cholesterol refeeding (80 µg/mL) following 10-h cholesterol deprivation. (E) Similar to D except the pre-experiment diet contained 1.2 µg/mL cholesterol. (F) Fat-body pS6 response during cholesterol re-stimulation (40 µg/mL) following 10-h feeding with 1.2 µg/mL cholesterol, in controls and fat-body *Tor* knockdowns. (G) Prothoracic-gland (PG) pS6 response during cholesterol re-feeding (40 µg/mL) following 1.2 µg/mL cholesterol limitation, in controls and with PG-specific *Tor* knockdown for 24 hours in the late third instar. Statistics: Data are shown as mean ±SEM, normalized to lowest-dose cholesterol or earliest re-feeding timepoint (cholesterol starvation, time 0). In (A, D, E, F, G, H), each point represents the cytoplasmic pS6 intensity of a single cell; in (B, C), each is a sample of multiple animals. (A, B, C, D, E, F, G) *p*-values calculated by Kruskal-Wallis ANOVA test with Dunn’s multiple comparisons.

When larvae were transferred from a cholesterol-depleted synthetic medium (0 µg/mL) to a cholesterol-replete one (80 µg/mL), we observed a time-dependent activation of the TOR pathway in the fat body (Figure 1D). Activation occurred within 30 minutes, peaking at 1 hour and subsequently exhibiting a dynamic oscillatory pattern. Taking into account the time required for food consumption and gut-mediated cholesterol distribution, cholesterol effects on TOR are rapid. Considering that natural food sources typically exhibit a range of sterol concentrations, rather than being absolutely sterol free, we investigated whether a transition from a low non-zero concentration to a cholesterol-replete diet would elicit a similar TOR-pathway activation pattern. Larvae were transferred to low-cholesterol food containing 1.2 µg/mL cholesterol for 10 hours at 84 hours AEL. In this condition, the transition to high (80 µg/mL) cholesterol at 94 h also resulted in increased fat-body pS6, but with levels peaking later at 6 hours post-shift (Figures 1D and 1E).

Developmental growth of larval tissues is modulated by the antagonistic actions of the growth-promoting, nutrient-responsive TOR and insulin pathways and the opposing effect of ecdysone signaling, which negatively regulates systemic growth via the receptor EcR^28^. Despite cholesterol’s being the precursor for steroid biosynthesis, we have previously demonstrated that cholesterol promotes growth in an ecdysone-independent manner by stimulating insulin secretion, a process dependent on the TOR pathway in the cells of the fat body and the blood-brain barrier ^11^. To substantiate this further, we compared the TOR-pathway response in the fat tissue of animals fed cholesterol to the response in those fed ecdysone, using concentrations of ecdysone sufficient to drive developmental progression^29^. Larvae were switched at 84 hours AEL from standard diet to a low (1.2 µg/mL)-cholesterol synthetic diet for 10 hours, after which they were placed on synthetic diet supplemented with either high cholesterol (80 µg/mL) or ecdysone. Dietary cholesterol induced a marked increase in fat-body pS6 levels compared to effects of dietary ecdysone, suggesting that cholesterol activates the TOR pathway through a mechanism that does not involve ecdysone (Figure S1A).

To confirm that the observed pS6 elevation in response to cholesterol availability was indeed mediated by the TOR pathway, we performed RNAi-mediated knockdown of *TOR* or its kinase substrate *S6K*, a key pathway component. Knockdown of *TOR* or *S6K* in the fat body (driven by *Cg-GAL4*, *Cg>*) abrogated the pS6 increase induced by dietary cholesterol (Figure 1F and S1B). We also examined the impact of dietary cholesterol on TOR activity in the prothoracic gland (PG), an endocrine tissue that requires substantial cholesterol flux for ecdysone synthesis and thus is particularly sensitive to cholesterol^24^. Cholesterol feeding increased pS6 levels in the PG, an effect that was dependent on TOR signaling, since it was abolished by PG-specific *TOR* knockdown. This was achieved by inducing *TOR* knockdown at 84 hours AEL [using temperature-sensitive (TS) *Tub-GAL80^TS^* with *phm-GAL4* – together, *phm^TS^>*], when larvae were transferred to low-cholesterol food containing 1.2 µg/mL cholesterol for 10 hours before refeeding with cholesterol-replete medium (40 µg/mL; Figure 1G). Additionally, employing a commercially available intermediate-sterol, cornmeal-based diet (NutriFly, NF)^11^, we demonstrated that larvae reared chronically on this diet supplemented with 40 µg/mL cholesterol throughout development exhibited elevated pS6 levels in the PG, indicative of enhanced TOR activity (Figure S1C). This effect was confirmed to be TOR-dependent, as the attenuation of TOR activity via knockdown eliminated the supplementation-induced increase in pS6 levels.

To further confirm the dynamic response of the TOR pathway to cholesterol, we used a transcriptional reporter of TOR function. In this system, expression of Luciferase from regulatory elements of the TOR-inhibited gene *unkempt* (*unk*) is inversely correlated with TOR activity^30^. When animals were transferred from a cholesterol-depleted synthetic medium (0 µg/mL) to a medium containing 80 µg/mL cholesterol, we observed a reduction of Luciferase activity in whole-larval lysates within 30 minutes, suggesting TOR-pathway activation (Figure 2A). This pattern mirrored the dynamics observed for the pS6 response (Figure 1D), and it was different from the systemic response to dietary protein refeeding following complete protein deprivation, which exhibited a gradual decline in reporter activity and thus indicates increasing TOR activity (Figure 2B).

**Figure 2:**
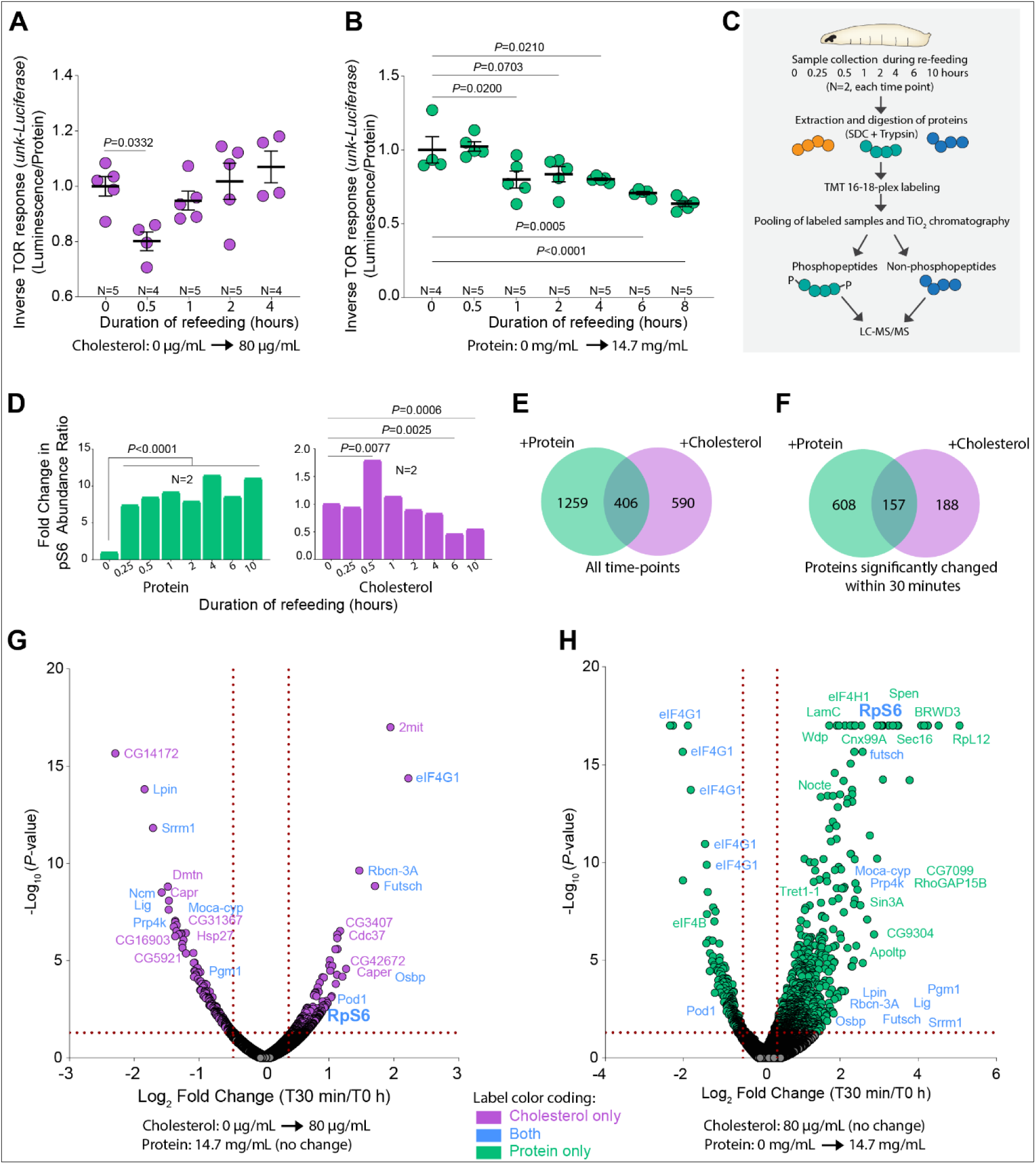
Cholesterol and protein feeding differently activate the TOR pathway and result in phosphorylation changes. (A) Whole-animal TOR response during to cholesterol refeeding (80 µg /mL), following 10-hour cholesterol deprivation, using *unkempt-Luciferase* assay. Lower values reflect greater Tor activity. (B) Whole-animal TOR response to protein refeeding (14.7 mg/mL casein) following 10-hour protein deprivation, using *unkempt-Luciferase* readout. Lower values reflect greater TOR activity. (C) Scheme of phosphoproteomic assays. (D) Mass-spectrometrically measured phosphorylation of ribosomal protein S6 on Ser239 (the phosphorylation site recognized by anti-pS6) over time in whole animals deprived of either protein (left) or cholesterol (right) for 10 hours then refed with complete medium. (E) Phosphoproteins significantly changed at any time point during refeeding with either protein or cholesterol – 406 phosphoproteins responded in both assays. (F) Phosphoproteins significantly changed after 30-minute refeeding with protein or cholesterol; 157 phosphoproteins responded in both assays. (G) Volcano plot of phosphoproteomic changes after 30 minutes’ cholesterol refeeding. See Figures S2A, S2C, S2E, S2G, S2I, and S2K for other time points. (H) Volcano plot of phosphoproteomic changes after 30 minutes’ protein refeeding. See Figure S2B, S2D, S2F, S2H, S2J, and S2L for further time points. Statistics: (A, B) Luciferase luminescence was normalized to the protein concentration of lysates. Mean ±SEM, normalized to deprivation condition (T0). Ordinary one-way ANOVA with Dunn’s multiple comparisons. (D, E, F) *p*-values were determined using the ANOVA test in Proteome Discoverer. (G, H) A phosphorylation change is considered significant with a 30% fold change (log_2_ fold change x>0.379 or x<-0.515) and *P*<0.05 (-log_10_ *P*>1.3).

We next employed an unbiased quantitative mass-spectrometry-based phosphoproteomics approach to further elucidate the signaling response to cholesterol (Figure 2C). These results show the same rapid and dynamic increase in S6 phosphorylation in response to cholesterol feeding when animals were transferred from a medium devoid of cholesterol (0 µg/mL) to one containing 80 µg/mL of this nutrient (Figure 2D, right panel). In contrast to this dynamic pattern, protein refeeding induced a sustained upregulation of TOR-pathway activity as reflected in pS6 levels (Figure 2D, left panel), recapitulating the results of the luciferase-reporter assay. A comparative analysis of proteins demonstrating significant phosphorylation changes (fold change upon transfer from cholesterol-or protein-starvation condition to a complete medium of greater than 30 percent, with a *p*-value less than 0. 05) at various refeeding intervals (15 minutes, 30 minutes, and 1, 2, 4, 6, and 10 hours) revealed 996 proteins that were (de)phosphorylated in response to dietary cholesterol, whereas 1,665 proteins showed altered phosphorylation in response to dietary protein (Figures 2E and 2F). Ribosomal protein S6 and eukaryotic Initiation Factor 4G1 (eIF4G1), another target of TOR that connects nutrient sensing to translation^31^, were among the proteins that were modified within 30 minutes in both cholesterol and amino-acid response (Figures 2G and 2H and S2A-S2L). This overlap indicates that TOR activity is rapidly activated by both dietary cholesterol and amino acids. The data also show dynamic changes in protein phosphorylation patterns over a 10-hour period when animals are refed cholesterol, a response phenomenon not exhibited after protein feeding. Together, these data show that dietary cholesterol feeding induces dynamic physiological responses, including TOR activation, in fat-body and PG cells and in whole animals. Furthermore, these changes are distinct in some ways from those that occur in response to amino-acid replenishment.

### HR3 is activated by cholesterol binding

Previous research suggests that HR3 interacts genetically with S6K^32^, and the closest human orthologs of HR3, RORα and -β, have been found to bind cholesterol^33^. However, the functional significance of this interaction remains unclear. The sequence similarity between the ligand-binding domains (LBDs) of HR3 and RORα^17^ suggests that cholesterol may also be a natural ligand of HR3. To determine whether HR3 binds cholesterol, we expressed the HR3 LBD in Hi5 insect cells and conducted affinity purification of this recombinant protein, followed by electrospray ionization mass-spectrometry (ESI-MS) under both denaturing and non-denaturing conditions to identify any co-purified ligands. In non-denaturing conditions, full-mass-range scans detected peaks suggesting a receptor:ligand complex with an added mass approximately equal to that of cholesterol (386.6 Daltons) (Figures 3A and 3B). To confirm the identity of the ligand, purified HR3 LBD was extracted with acidified chloroform/methanol to isolate co-purifying small molecules, which were derivatized and analyzed by electron ionization gas chromatography-mass spectrometry (EI-GC/MS). Chromatography of the extract yielded a single major peak at 19 minutes (Figure 3C) which, when ionized, produced a spectrum characteristic of derivatized cholesterol (Figure 3D). A cholesterol standard processed through the same protocol yielded a close match to the spectrum of the putative ligand (Figure 3E and 3F). Taken together, evidence from the combination of protein ESI-MS and GC/MS demonstrates that the LBD of HR3 binds cholesterol with a 1:1 stoichiometry.

**Figure 3:**
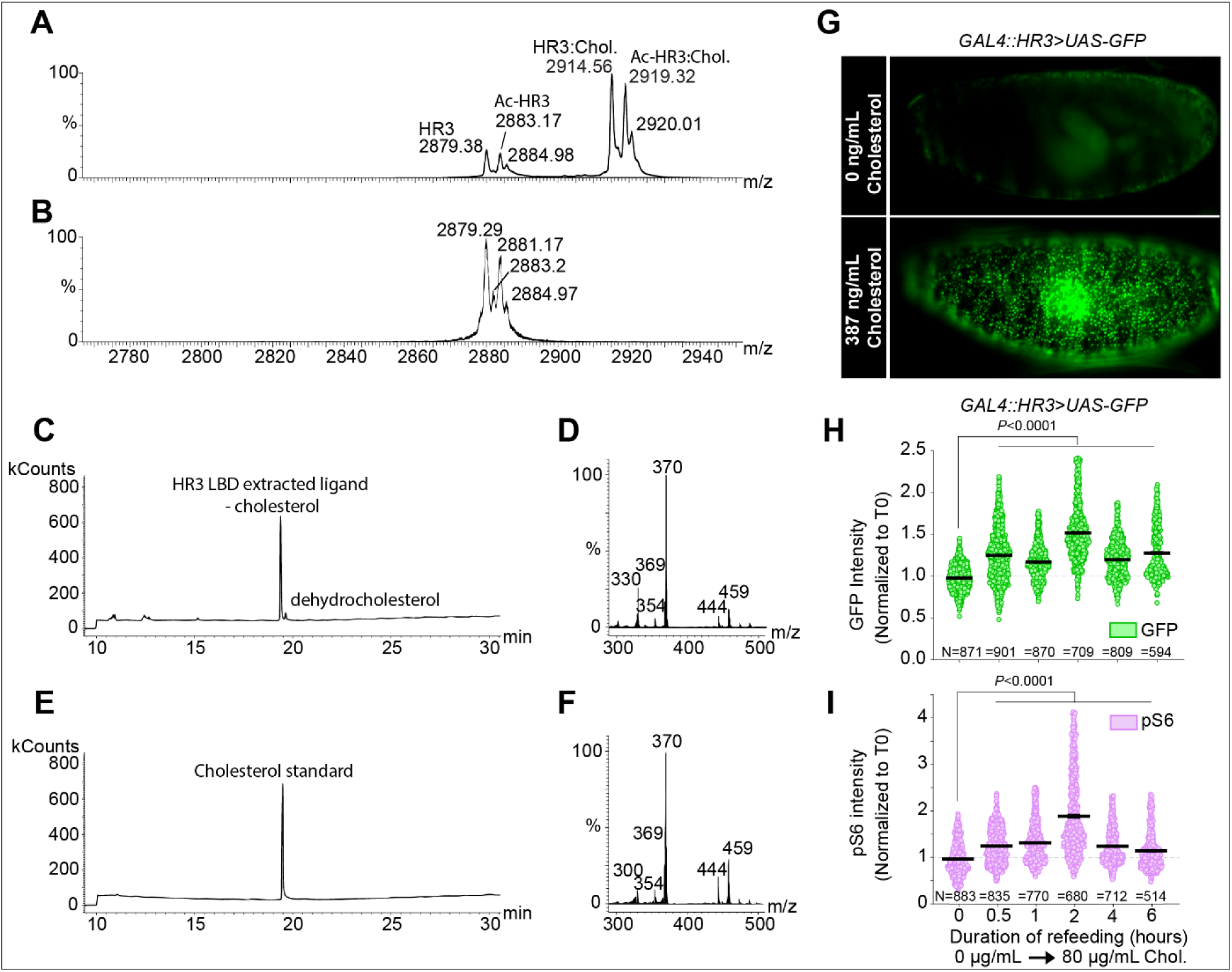
Cholesterol is a ligand of HR3. (A) Mass spectrum of purified HR3 under non-denaturing conditions with peaks representing HR3 (2879.38 *m/z*), acetylated HR3 (2883.17 *m/z*, Ac-HR3), liganded HR3:cholesterol complex (2914.56 *m/z*, HR3:Chol.), and acetylated HR3:cholesterol complex (2919.32 *m/z*, Ac-HR3:Chol.) indicated. (B) Quadrupole time-of-flight collision-induced dissociation MS spectrum of purified HR3 after the loss of the ligand. (C) A gas chromatogram of a derivatized chloroform/methanol extraction of HR3 LBD produces a peak at the indicated time representing the non-ionizing ligand of purified HR3 LBD. (D) The corresponding electron ionization (EI) spectrum of the fraction containing the major peak. (E) Gas chromatogram of a derivatized cholesterol standard. (F) The EI fragment spectrum from the peak fraction containing the standard. (G) GFP fluorescence was recorded from permeabilized living embryos expressing the HR3 ligand sensor *GAL4::HR3>UAS-GFP,* cultured for 1 hour either in the absence of cholesterol (top) or with 387 ng/mL (1 µM) cholesterol in the medium (bottom). (H, I) *Drosophila* larval fat-body GFP signals of *GAL4::HR3>UAS-GFP* (H) and anti-pS6 in the same animals (I) in response to cholesterol re-feeding (80 µg/mL) following 10 hours’ cholesterol deprivation. Statistics: (H and I) Data are shown as means ±SEM, normalized to T0. Each data point represents a single cell measurement. Significance determined by Kruskal-Wallis nonparametric ANOVA with Dunn’s multiple comparisons.

To assess the binding of cholesterol to HR3 *in vivo*, along with functional outcomes of this binding, we tested the ability of cholesterol to activate a transgenic HR3 ligand-sensor construct (GAL4::HR3). In this system, the LBD of HR3 is fused with the DNA-binding domain (DBD) of GAL4. Ligand binding to the LBD of the fusion protein activates expression of a *UAS-GFP* reporter gene^34^. Cholesterol addition to permeabilized living embryos carrying this system induced widespread GFP expression, indicating that exogenous cholesterol can bind and activate the LBD of HR3 (Figure 3G).

We next used this ligand sensor to examine whether the LBD of HR3 can be activated by cholesterol derived from the diet. Cholesterol refeeding of larvae after a deprivation period on synthetic diet lacking cholesterol led to increased GFP staining in the fat body, consistent with binding of cholesterol (or a derivative) obtained through the diet (Figure 3H). We also confirmed that this cholesterol treatment produces the characteristically dynamic fat-body TOR activation pattern as measured by pS6 staining (Figure 3I). These findings collectively indicate that cholesterol binds to the LBD of HR3 and that this protein functions as a cholesterol-responsive receptor.

### HR3 modulates TOR activity and regulates systemic growth

Given the activation of HR3 by dietary cholesterol, we investigated whether HR3 might influence systemic growth via modulation of the TOR signaling pathway on standard diet. Knockdown of *HR3* in the fat body or PG using an RNAi construct that targets all of the annotated transcript variants of *HR3* led to reductions in pS6 staining in these tissues (Figures 4A and 4B). This attenuation indicates a regulatory interaction whereby HR3 contributes to S6 phosphorylation, perhaps mediated by TOR and S6K. To more directly show that phosphorylation of S6K and S6 occurs downstream of HR3, we simultaneously expressed a constitutively active S6K allele in the fat body, which suppressed the pS6 reduction induced by *HR3* knockdown (Figure 4C), implying that HR3 modulates S6 phosphorylation via the TOR-S6K signaling pathway. It is well established that reduced fat-body TOR activity attenuates systemic growth^35^. In line with this, we found that *HR3* knockdown in the fat body reduced systemic growth, as reflected in a decrease in larval mass, validated using two distinct fat-body-specific drivers (*Cg>* and *pumpless-GAL4*, *ppl>*) and by three independent RNAi lines (Figures 4D and 4E and S3A).

**Figure 4:**
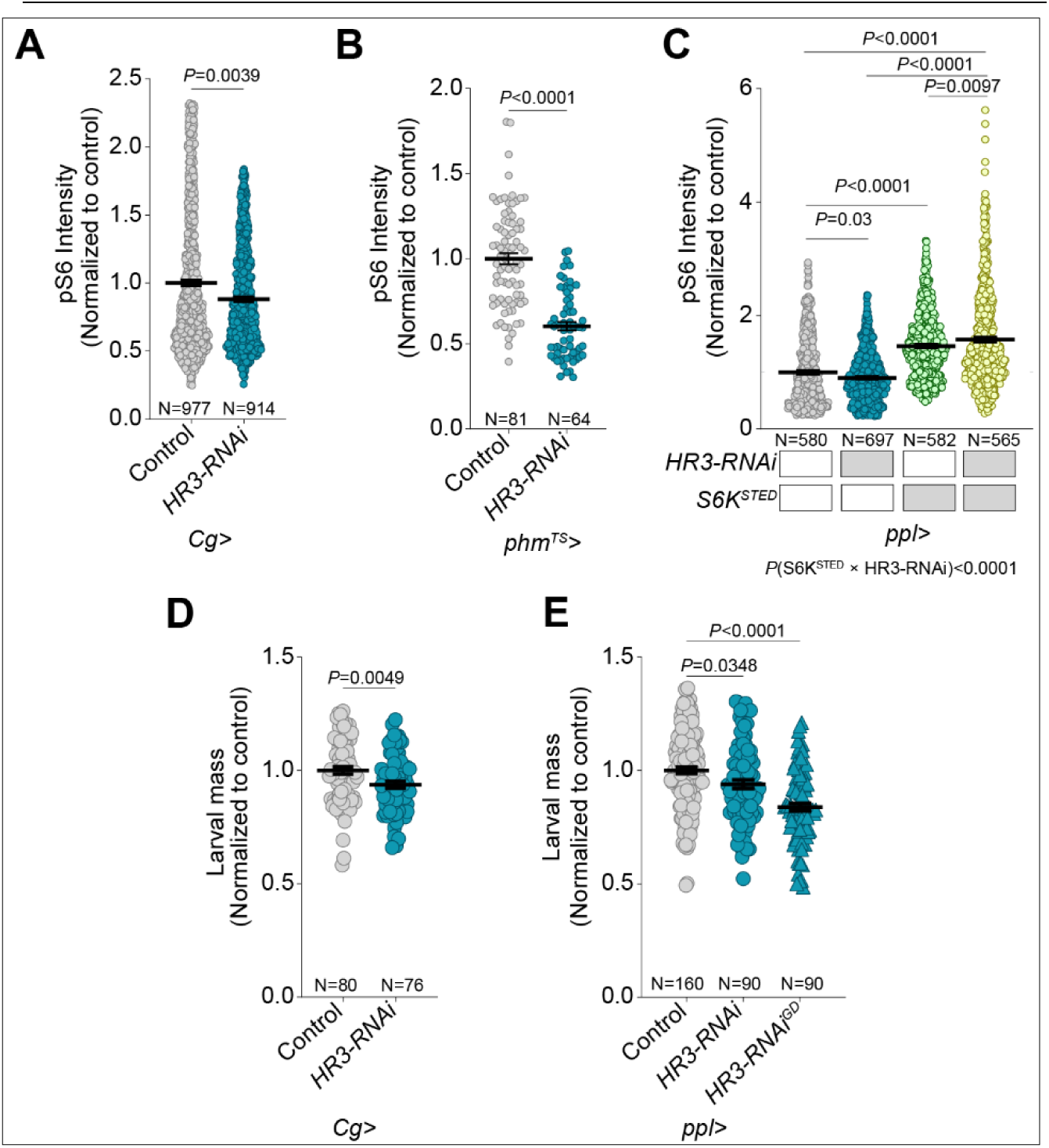
Knock-down of *HR3* decreases systemic growth and TOR activity. (A) *Drosophila* fat-body pS6 quantification in fat-body cells of controls and animals expressing RNAi against all isoforms of *HR3* in the fat body using *Cg-GAL4*. (B) Quantification of pS6 in the cells of the PG in controls and animals expressing *HR3* knockdown using *phm^TS^*. (C) Quantification of pS6 in fat-body cells of control larvae and animals expressing *HR3* RNAi in the fat tissue using *ppl>*, with and without co-expression of constitutively active S6K (*S6K^STED^*). (D) Body mass of control larvae and larvae expressing fat-body knock-down of *HR3* using *Cg-GAL4*. (E) Body mass of control larvae and animals expressing either of two independent HR3-RNAi constructs using a second driver (*ppl-GAL4*). Statistics: (A, B, C, D, E) Graphs plot mean ±SEM. (A, B, D) Significance determined using two-tailed unpaired t-test (C, E) Kruskal-Wallis ANOVA test with Dunn’s multiple comparisons, with two-way ANOVA test for epistasis in C. In (A, B, C) each data point reflects a single cell; in (D, E) each point reflects a single larva.

### HR3 regulates TOR pathway activity in response to cholesterol

Given the role of HR3 in binding cholesterol and modulating TOR pathway activity, we investigated whether HR3 is necessary for the cholesterol-induced activation of the TOR pathway. Larvae, at 84 hours AEL, were first placed on a 1.2-µg/mL-cholesterol diet for 10 hours, followed by a shift to a high-cholesterol diet (80 µg/mL). Fat-body-specific knockdown of all *HR3* variants through RNAi resulted in diminished cholesterol-induced activation of the TOR pathway over 6 hours, as indicated by reduced pS6 levels (Figure 5A). Consistent with our previous findings (Figure 1E), we found that elevated dietary cholesterol rapidly activated TOR signaling in fat tissue within 1 hour, with the activation progressively intensifying up to 6 hours post-transfer. By contrast, cholesterol replenishment failed to elevate pS6 levels within an hour in animals with fat body-specific *HR3* knockdown, and the long-term activation was attenuated in these animals. These findings were further corroborated using two further independent RNAi constructs also targeting all *HR3* isoforms, showing that fat-body-specific knockdown of *HR3* largely abolished cholesterol-stimulated TOR pathway activation (Figure 5B). We next explored whether HR3 selectively activates TOR in response to cholesterol. Contrary to the diminished TOR activation by cholesterol, the response of TOR to dietary amino acids remained unperturbed by *HR3* knockdown (Figure 5C). This indicates that HR3 does not influence the activation of TOR by dietary amino acids, distinguishing its role in cholesterol-mediated TOR activation.

**Figure 5:**
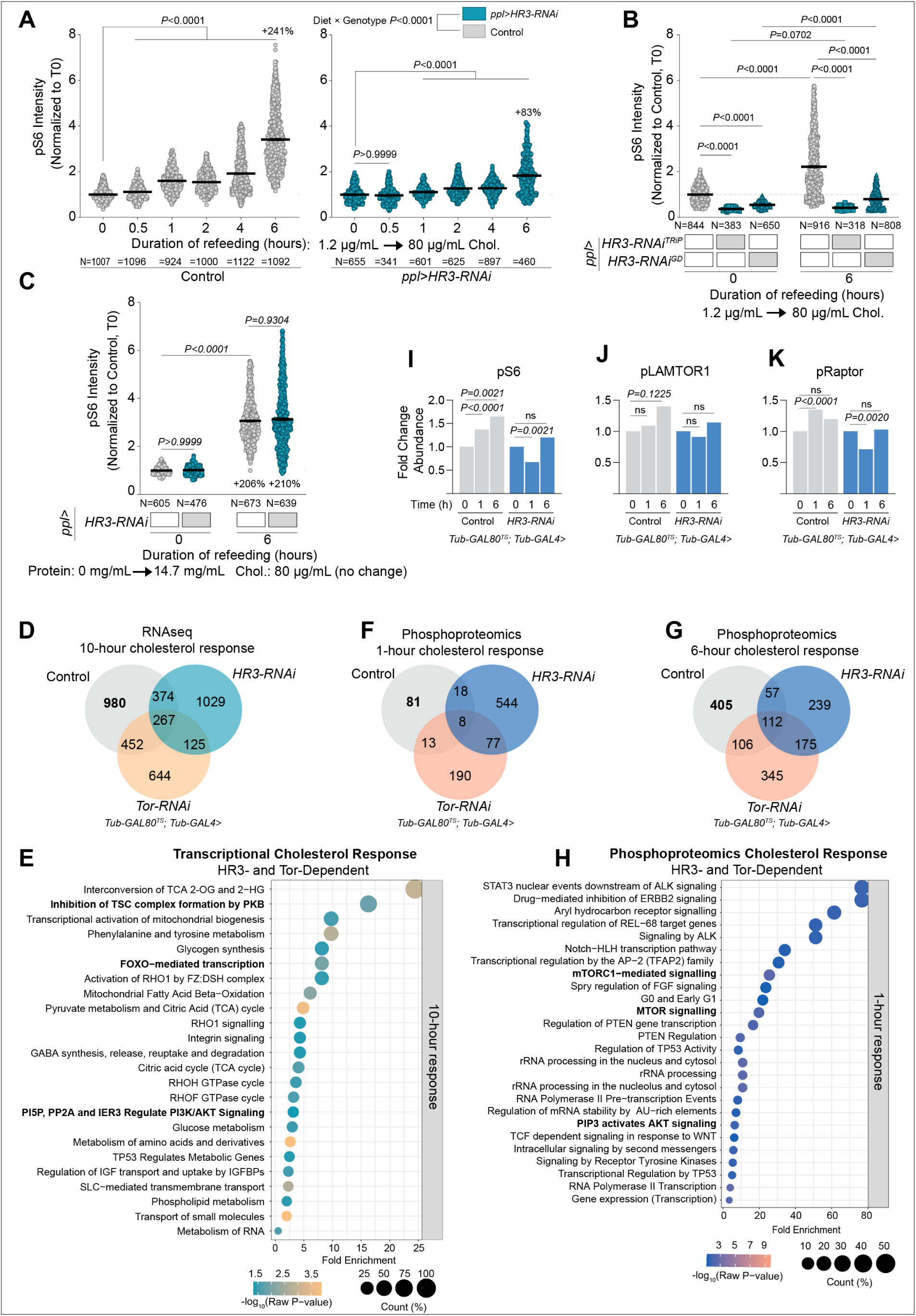
HR3- and Tor-dependent cholesterol response. (A) Phospho-S6 levels in larval fat-body cells in controls and animals expressing *HR3* RNAi in the fat body using *ppl-GAL4*, measured over time after cholesterol refeeding (80 µg/mL) following 10 hours on low- (1.2 µg/mL)-cholesterol medium. (B) pS6 levels in larval fat-body cells in controls and animals expressing two independent *HR3-RNAi* constructs, measured six hours after refeeding with 80 µg/mL cholesterol following 10 hours’ feeding with low-cholesterol medium. (C) pS6 levels were measured in larval fat-body cells of controls and animals with fat-body specific *HR3* knockdown using *ppl-GAL4*. Measurements were taken after amino-acid refeeding (14.7 mg/mL) following 10 hours on protein-free medium. (D-K) Transcriptional and proteomic responses to cholesterol refeeding (80 µg/mL) over time in control *Tub-GAL80^TS^; Tub-GAL4* larvae and those ubiquitously expressing *HR3-RNAi* or *TOR-RNAi,* following 10-hour low-cholesterol (1.2 µg/mL) feeding. (D) Number of genes differentially expressed between the three genotypes at the 10-hour time point. In bold, “980” reflects genes whose cholesterol response is dependent on both HR3 and TOR. (E, H) Selected pathways enriched in these 980 genes, identified using Panther and Reactome. (F, G) Significant phosphorylation changes in response to cholesterol feeding after 1 and 6 hours. Numbers in bold (top left segment of each diagram) represent changes that require both HR3 and TOR. (I, J, K) Phosphorylation of S6, LAMTOR1 or RAPTOR over time in response to cholesterol re-feeding in control and HR3-knockdown animals. Statistics: (A-C) Graphs plot mean ±SEM, each data point representing a single cell. Data normalized to control at T0. Statistical significance determined using Kruskal-Wallis ANOVA test with Dunn’s multiple comparisons. Interaction between diet and genotype was determined using two-way ANOVA. (D) For RNA-seq data model-dependent *p*-values were used. (F, G) Changes greater than 30% in magnitude with *p*<0.05 are taken as significant. *p*-values were determined using the ANOVA test in Proteome Discoverer. (E, H) pathway analysis with Fisher’s t-test; only pathways with *p*<0.05 are included. (I, J, K) *p*-values were determined using ANOVA test in Proteome Discoverer.

To further examine any potential genetic interaction between HR3 and EcR, we knocked down *EcR* in the fat body and assessed the TOR pathway’s response to cholesterol. Given that HR3 is an ecdysone-inducible gene^17^, loss of *EcR* would be expected to reduce *HR3* expression, which would in turn dampen the TOR response to cholesterol, similar to the effects of HR3 knockdown itself. Contrary to this expectation, *EcR* knockdown in the fat body enhanced the response to cholesterol (Figure S3B). EcR suppresses cellular cholesterol uptake^36^, so this result is consistent with increased cholesterol uptake that activates TOR. The opposite effects of *HR3* and *EcR* knockdowns suggest that HR3 contributes to cholesterol-induced TOR activation in an EcR-independent manner.

To gain deeper insight into the HR3-mediated mechanisms underlying cholesterol-induced TOR activation, we profiled the genomic and phosphoproteomic responses to cholesterol in animals ubiquitously expressing knockdown of either *HR3* or *TOR*. Using *Tub-GAL80^TS^* in conjunction with the *Tub-GAL4* driver, RNAi was induced in larvae at 111 hours AEL by switching animals from 18 °C to 29 °C. Nine hours later, these animals were switched from a standard diet to a low-cholesterol diet (1.2 µg/mL) for 10 hours. Subsequently, larvae were transferred to a high-cholesterol diet (80 µg/mL) for varying lengths of time to assess their response. In our RNA-seq profiling efforts, we identified 2073 genes exhibiting significant regulatory changes (both upregulation and downregulation) in response to a high-cholesterol diet (80 µg/mL) in control larvae, compared against age-matched larvae maintained on a low cholesterol diet (1.2 µg/mL), which served as reference point for all comparisons (Figure 5D). The transcriptional response to cholesterol was strongly dependent on both HR3 and TOR, as evidenced by a loss of cholesterol response in 980 (47%) of these genes in animals lacking either *HR3* or *TOR*. This underscores the critical roles of HR3 and TOR in orchestrating the transcriptional machinery responsive to cholesterol. Additionally, other sets of genes became aberrantly responsive to cholesterol upon the knockdown of either HR3 or TOR, with an overlap of 125 genes between these sets, suggesting that these factors normally act to repress such responses. Collectively, these data imply that HR3 and TOR are required to initiate a specific transcriptional response to cholesterol through regulatory mechanisms that activate certain genes while suppressing the response of others to cholesterol. Further investigation was carried out through Reactome pathway analysis applied to the 980 genes that require both HR3 and TOR for proper cholesterol response. This analysis revealed a significant enrichment of genes involved in pathways that include the TSC2 complex and insulin signaling, specifically PI3K/AKT/FOXO-related pathways (Figure 5E). Given that TSC2 is a critical inhibitor of the TOR signaling pathway, and considering the interplay between TOR and insulin signaling, these findings further support our model in which cholesterol modulates TOR signaling via a mechanism that requires HR3.

We next conducted phosphoproteomic analysis for deeper insights into the HR3 and TOR-dependent pathway responses to cholesterol. All comparisons were made against the baseline established at the zero-hour-refeeding time point of control larvae maintained on a 1.2-µg/mL cholesterol diet for the preceding 10 hours, before transfer to cholesterol-replete medium. Our results indicated a rapid (within 1 hour) alteration of phosphorylation of 81 proteins (those showing a change of over 30 percent with a *p*-value of less than 0.05) that are HR3- and TOR-dependent (they do not occur in the absence of either protein; Figure 5F). Notably, a substantial number of proteins (544) changed phosphorylation in response to cholesterol in *HR3* knockdown animals but not in controls, suggesting that HR3 normally suppresses these changes. Mirroring the pattern seen in fat-body pS6 levels upon transferring animals from 1.2-µg/mL to 80-µg/mL cholesterol (Figures 1E and 5A), the cholesterol response was greater at 6 hours relative to 1 hour, with 405 proteins exhibiting HR3-and- TOR-dependent phosphorylation changes (Figure 5G). Pathway analysis of this HR3-and-TOR-dependent phosphoproteomic response highlighted the TOR pathway, along with ribosome- and translation-related processes, which are regulated by TOR activation (Figures 5H and S3C). The HR3-dependent phosphorylations included S6 itself (Figure 5I), consistent with the HR3-dependent cholesterol-mediated phosphorylation of S6 in the fat body (Figures 5A and 5B), as well as LAMTOR and RAPTOR (Figures 5J and 5K), two key regulatory components acting upstream of TOR that regulate its activity via interactions with the Rags. This implies HR3 operates at the level of, or upstream of, LAMTOR and RAPTOR to modulate TOR activity, consistent with insights from other recent studies in mammalian systems^3,37^. At the 6-hour time point, only 239 cholesterol-induced changes in protein phosphorylation were observed in *HR3* knockdown animals (Figure 5G), in contrast to the 544 alterations observed at the one-hour time point (Figure 5F). This pattern suggests that while HR3 normally activates the TOR pathway within 1 hour in response to cholesterol, it concurrently modulates a broad spectrum of proteins and blocks their phosphorylation, presumably to maintain a balanced cellular response. This dual role of HR3 suggests a complex regulatory mechanism through which it both stimulates and restrains the phosphorylation of proteins in a time-dependent manner in response to cholesterol levels, ensuring a precisely controlled cellular adaptation.

### Cholesterol-induced TOR activation via HR3 is independent of its DNA-binding function

To delineate how HR3 conveys cholesterol availability to the TOR signaling cascade, we investigated the hierarchical relationship between HR3 and TOR. Given the previously documented genetic interaction between S6K and an HR3 isoform lacking the DNA-binding domain (DBD)^32^, we explored whether this truncated isoform participates in HR3’s mediation of cholesterol’s physiological effects (Figure 6A). Overexpression of this shorter isoform – HR3(LBD) – in the fat body led to increased systemic growth, as evidenced by increased larval weight (Figure 6B). To ascertain whether the systemic growth effects of HR3(LBD) expression were mediated by the TOR pathway, we simultaneously silenced TOR expression in the fat body, which in turn abrogated the growth-promoting effects of the overexpressed HR3(LBD) variant (Figure 6C). This finding indicates that the truncated HR3 isoform requires TOR function for its growth-promoting effects and suggests that HR3 can modulate TOR-pathway activity through a mechanism that does not require its DBD function. To pursue this notion further, we assessed whether HR3(LBD) is a mediator of cholesterol-stimulated TOR pathway activation. Chronic exposure to a cholesterol-enriched diet of NF supplemented with 40 µg/mL cholesterol combined with overexpression of HR3(LBD) in the fat body significantly enhanced pS6 levels, suggesting that HR3(LBD) promotes cholesterol-induced TOR-pathway activation (Figure 6D). Aligning with phosphoproteomics findings that potentially position HR3 upstream of the Rags, the augmented pS6 levels caused by HR3(LBD) overexpression were entirely negated by concurrent *TOR* knockdown, demonstrating that HR3(LBD) acts at or above the level of TOR.

**Figure 6:**
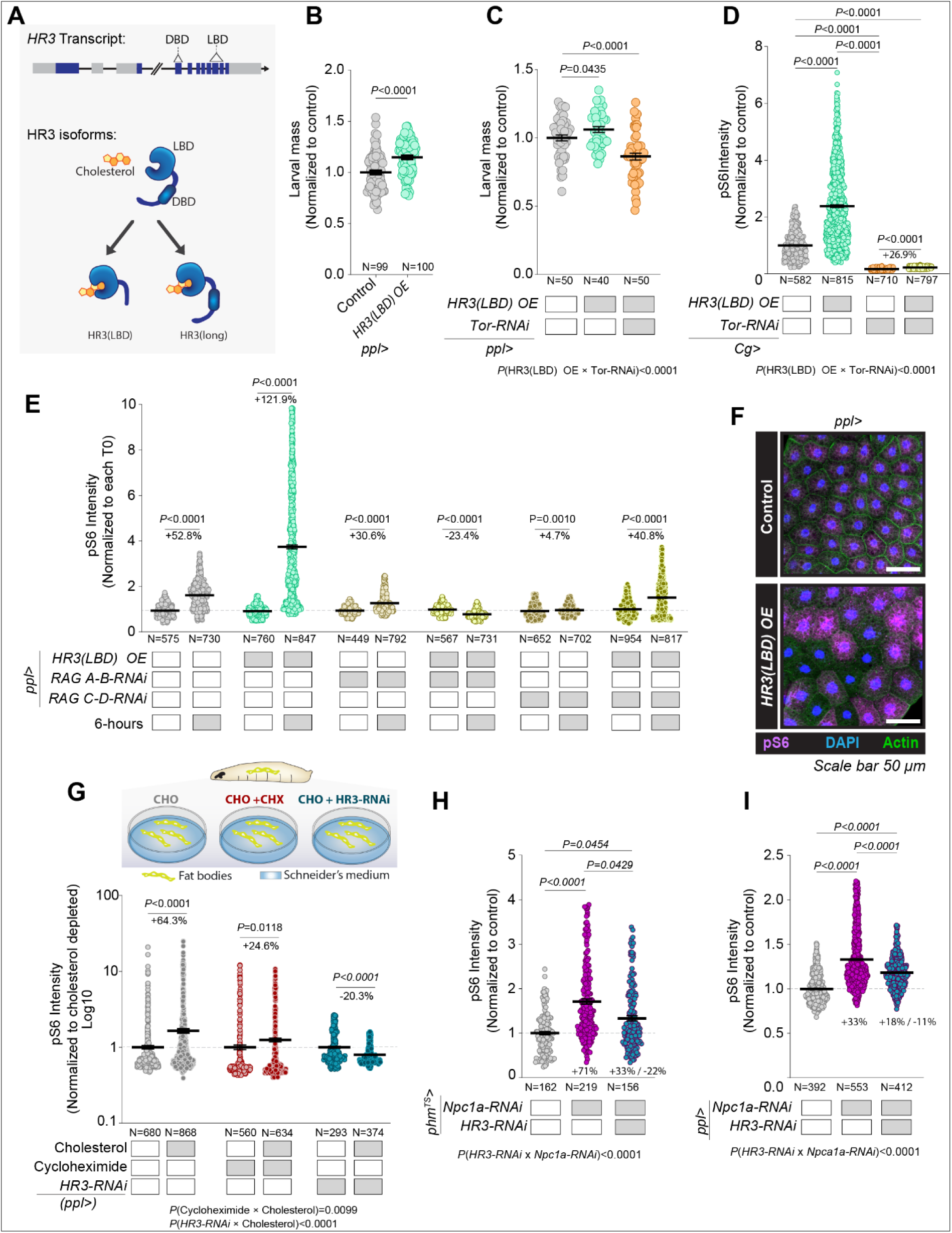
HR3 mechanism of TOR pathway modulation. (A) (Top) Representative illustration of the *HR3* transcript which can be alternatively spliced to form at least 7 different isoforms. The DNA binding domain (DBD) is found in the 3^rd^ exon, while the ligand-binding domain spans exons 7-9. Exons shown in blue, introns in grey. A large intronic region is removed and represented with a line break. (Below) Representation of two major types of isoforms, the major one containing both a DBD and a LBD [called HR3(long)] and an alternative isoform without a DBD [called HR3(LBD)], a novel transcript not yet annotated in FlyBase that lacks a first alternative exon^32^. The *HR3* knock-down RNAi lines targets all known isoforms, whereas the HR3(LBD) over-expression (OE) construct only increases expression of HR3(LBD). (B) Larval mass at 96 h AEL of controls and animals overexpressing *HR3(LBD)* in the fat body using *ppl>*, grown on STD-FF. (C) Larval mass of controls and animals overexpressing *HR3(LBD)* in the fat body (*ppl*>) with and without simultaneous *Tor* knockdown, grown on STD-FF. (D) Phospho-S6 staining in 96-hour-AEL animals with and without fat-body-specific HR3(LBD) overexpression, with and without simultaneous *Tor* knockdown, chronically fed a high-cholesterol diet (NF+40 µg/mL). (E) Phospho-S6 staining in fat-body cells of controls and animals overexpressing HR3(LBD) in the fat body, with and without simultaneous knockdown of *RagA-B* or *RagC-D*, with and without 6-hour cholesterol refeeding (80 µg/mL) following 10-hour feeding on low-cholesterol medium (1.2 µg/mL). (F) Representative pS6 stains of fat-body tissue from controls and animals overexpressing HR3(LBD) in the fat body using *ppl>*, after 6 hours’ cholesterol re-feeding (80 µg/mL). Scale bars, 50 µm. Figures S4C and S4C quantify the variability in pS6 staining. (G) Fat-body tissue dissected from 96-h-AEL larvae (controls and fat-body-specific *HR3* knockdowns) was cholesterol-depleted *ex vivo* with MCD over 2 hours and replenished using MCD:cholesterol over 1 hour, with and without cycloheximide in the medium. (H) pS6 staining intensity in cells of the PG in controls, animals expressing *Npc1a* RNAi, and in animals expressing RNAi against *Npc1a* and *HR3*. (I) pS6 staining intensity in cells of the fat body in controls, animals expressing *Npc1a* RNAi, and in animals expressing RNAi against *Npc1a* and *HR3*. Statistics: Data are normalized to controls and graphed as mean ± SEM. (E,B,G) Each data point represents a single cell. Data pairs are normalized to each cholesterol-starvation condition or controls. Pairwise *P*-values were calculated using Mann-Whitney tests. Significant interactions between variables were assessed by two-way ANOVA. (C, D, H, I) Each data point represents a single cell. Kruskal-Wallis ANOVA with Dunn’s multiple comparisons (C, D, G, H, I) Significant interactions between variables were assessed by two-way ANOVA.

Next, we asked whether HR3(LBD) is involved in the acute activation of the TOR pathway in response to cholesterol. Larvae, 84 hours AEL, were initially fed a diet containing 1.2 µg/mL cholesterol for 10 hours, then transferred to a diet with the higher cholesterol concentration of 80 µg/mL to examine their response. In animals overexpressing HR3(LBD) in the fat body and fed 80 µg/mL cholesterol for 1 hour, pS6 levels were significantly elevated compared to controls, an effect that was blocked by TOR knockdown (Figure S4A). The increase in pS6 was even more pronounced and substantial at the 6-hour time point, indicating that the HR3(LBD) isoform is a key mediator of cholesterol-induced TOR-pathway activation (Figure 6E). This effect was not produced when the diet was supplemented with ecdysone, instead of cholesterol, demonstrating that HR3 mediates a cholesterol-specific response (Figure S4B). Notably, this HR3(LBD) overexpression also induced significant intercellular variability in pS6 levels after 6 hours of cholesterol stimulation (Figures 6F, S4C and S4D), suggesting that overexpressing *HR3(LBD)* leads to a progressive destabilization or desynchronization of TOR-pathway dynamics. These findings support the hypothesis that HR3 plays a cell-autonomous role in cholesterol sensing, acting upstream of TOR.

### HR3 mediates cholesterol-responsive TOR signaling via genomic and non-genomic mechanisms upstream of the Rag GTPases

Nuclear receptors are ligand-regulated transcription factors that canonically exert their effects through transcriptional regulation ^17^. However, rapid effects are often mediated by non-genomic actions in which the receptor is localized in membranes or the cytosol and directly influences signal-transduction pathways^38,39^. Given that HR3(LBD) mediates rapid non-genomic effects^32^, we asked whether cholesterol-mediated TOR activation might entail both genomic and non-genomic HR3 functions. To determine the mechanism by which cholesterol activates TOR, we inhibited translation using cycloheximide to suppress any regulation of the TOR pathway by newly expressed proteins. We cultured *ex-vivo* fat-body tissues from larvae at 96 AEL in Schneider’s medium supplemented with lipid-depleted serum. Tissues were depleted of sterols through a 2-hour incubation with methyl-beta-cyclodextrin (MCD, 0.75%), after which they were stimulated with cholesterol complexed with MCD (0.1%). This *ex-vivo* cholesterol stimulation increased fat-body pS6 levels in tissues dissected from control animals (Figures 6G and S4E). Cholesterol stimulation also increased pS6 levels in fat-body tissues treated with cycloheximide, although not as robustly as in controls without cycloheximide. This suggests that a significant portion of the rapid response does not require new protein synthesis. Together this implies that cholesterol activates the TOR pathway through mechanisms independent of translation, and potentially transcription, as well as through processes that require new protein synthesis and could involve a transcriptional response. We then assessed whether HR3 is integral to both processes by examining the pS6 response to cholesterol in fat bodies in which HR3 expression had been silenced. Fat bodies lacking *HR3* exhibited no upregulation of pS6 in response to *ex-vivo* cholesterol stimulation (Figure 6G), indicating that HR3 is essential for TOR-pathway activation by cholesterol. This demonstrates that cholesterol activates the TOR pathway through HR3 via tissue-autonomous mechanisms that may involve transcriptional regulation as well as non-genomic actions, such as interacting with signaling proteins upstream of TOR, such as the Rag GTPases. We therefore assessed the intracellular localization of HR3 and discovered that this nuclear receptor is not restricted to the nucleus, but also exhibits a cytosolic or membrane localization pattern (Figure S4F). This observation supports the possibility that HR3 may interact with components of the TOR pathway in the cytosol or at the surface of lysosomes. To further this, we explored whether inhibiting the Ragulator-Rag complex would negate the effects of HR3(LBD) overexpression. The TOR activity increase due to HR3(LBD) overexpression was largely dependent on both RagA-B and RagC-D (Figure 6E). We then induced lysosomal cholesterol accumulation by depleting *Npc1a*, which resulted in TOR activation. By simultaneously reducing *HR3*, we observed a decrease in the hyperactivation of TOR caused by *Npc1a* knockdown (Figures 6H, 6I and S4G). This indicates that HR3 is required for TOR activation in response to cholesterol accumulation within lysosomes. Collectively, our findings support a model in which HR3, upon binding cholesterol, intersects the TOR pathway at the level of, or upstream of, Rag-mediated TOR activation, similar to amino-acid mediated TOR activation, which occurs through effects on the Rag proteins induced by a diverse collection of sensory proteins.

### Human RORα activates the TOR pathway in response to exogenous cholesterol

Considering the functional and sequence conservation between human RORα and its *Drosophila* counterpart, HR3, we investigated the potential role of RORα in modulating mammalian TOR-pathway activity in response to cholesterol. We used human KARPAS-707H cells, a multiple myeloma cancer cell line strongly expressing RORα, to assess TOR signaling in response to exogenous cholesterol. *In-vitro* cholesterol depletion (0.75% MCD for 2 hours) of these cells reduced TOR signaling, reflected in decreased pS6 abundance, whereas cholesterol repletion (complexed with 0.1% MCD) enhanced pS6 levels dose-dependently (Figures 7A and 7B), establishing a direct correlation between extracellular cholesterol levels and TOR-pathway activation. *RORα* knockdown using synthetic siRNA attenuated the pS6 response to cholesterol (Figure 7C), mirroring the *HR3*-knockdown effect observed in *Drosophila* (Figures 5A and 5B).

**Figure 7:**
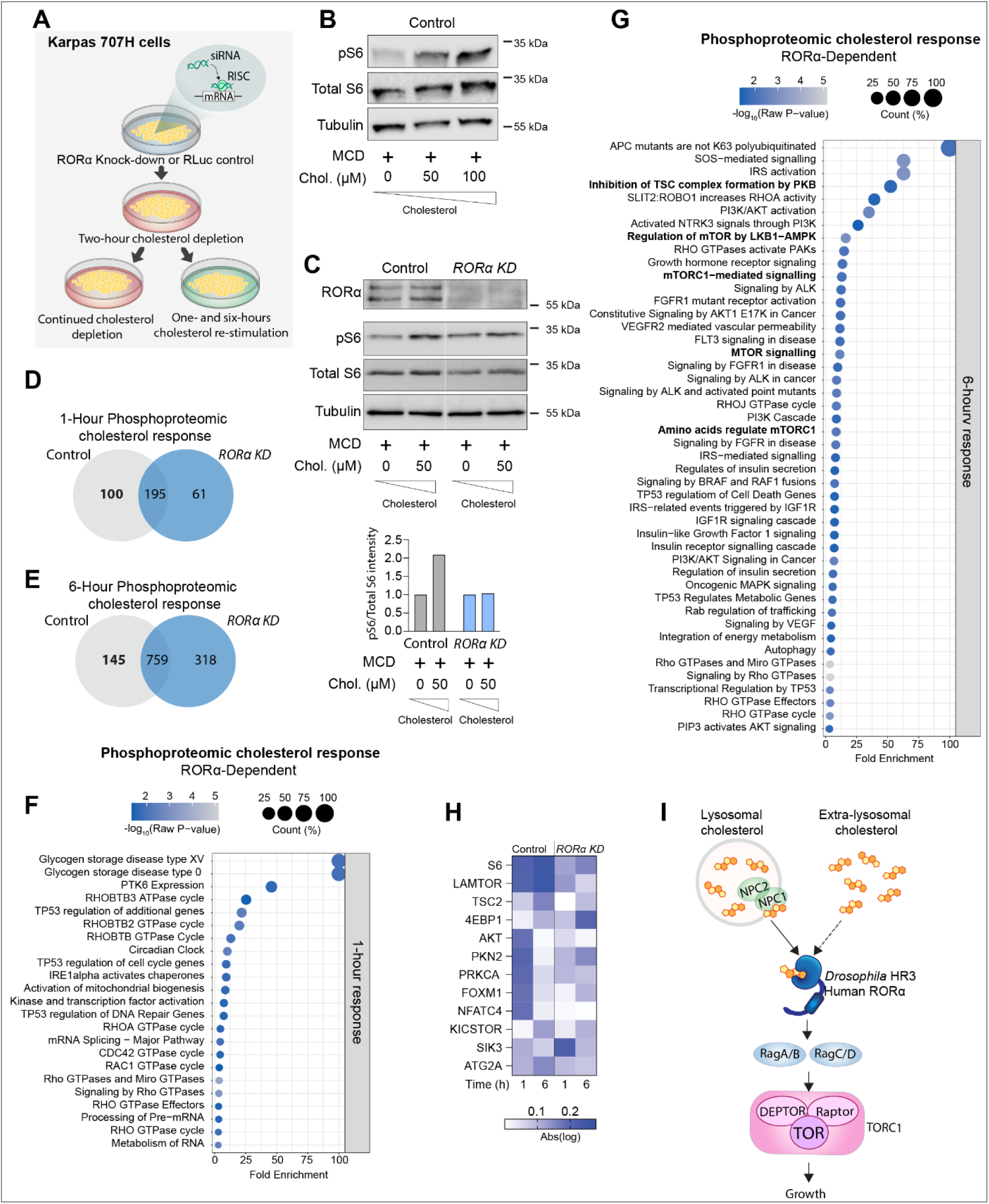
Conserved mechanism of cholesterol regulation of the TOR pathway by human RORα *in vitro*. (A) Illustration of experimental treatments of *in vitro* experiments using human multiple myeloma cell line Karpas 707H, chosen for its high expression of *RORα*. *RORα* was knocked down in the cells using siRNA, and the cells underwent a two-hour cholesterol-depletion treatment using 0.75% methyl-β-cyclodextrin (MCD) and 0.5% lipid-depleted serum (LDS). The cells were replenished with cholesterol using MCD:cholesterol (0.1% MCD:50 µM cholesterol or 0.2% MCD:100 µM cholesterol) with 0.5% LDS or simply changed to new depletion medium and were sampled 1 and 6 hours later. Protocol based on Shin et al.^37^ and Castellano et al.^3^ (B) Immunoblot against pS6, total S6, and α-Tubulin of control and KARPAS-cell extracts after 1-hour cholesterol replenishment or mock treatment. (C) Immunoblot against RORα, pS6, total S6, and alpha-Tubulin of extracts from control and RORα-siRNA cells, following 1-hours cholesterol replenishment or mock treatment. The ratio of pS6 to total S6 staining is quantified, normalized to cholesterol-depleted condition for each genotype. (D, E) Phosphoproteomic differences between cholesterol-replenished and mock-treated cholesterol-depleted control and RORα-siRNA-treated cells after 1 (d) and 6 (e) hours of treatment. The bolded control number reflects phosphorylation changes that require RORα. (F, G) Selected pathways enriched in proteins exhibiting *RORα*-dependent cholesterol response identified using Panther and Reactome. (H) RORα-dependent phosphorylation response of TOR and insulin related proteins in response to re-stimulation of cholesterol within 1 or 6 hours. (I) Model proposed for the activation of TOR signaling mediated by HR3 and RORα in response to cholesterol. Dashed line indicates a possible non-lysosomal pool of cholesterol activating HR3/ RORα. Statistics: (D, E) Changes greater than 30% up or down with *p*<0.05 were considered significant. *p*-values were determined using the ANOVA test in Proteome Discoverer. (F, G) Pathway analysis with Fisher’s T-test; only pathways with p<0.05 are included.

For more comprehensive insights into RORα-mediated TOR-pathway activation by cholesterol, we performed phosphoproteomic analysis of KARPAS-707H cells. Comparisons were made against cholesterol-depleted cells, considering a 30% change in phosphorylation with a *p*-value<0.05 to be significant. Within 1 hour of cholesterol replenishment, 100 significant RORα-dependent protein phosphorylation changes occurred, increasing to 145 changes after 6 hours (Figures 7D and 7E). Pathway analysis showed a pronounced enrichment in components of the TOR and PI3K/AKT pathways among the RORα-dependent cholesterol-induced phosphorylation changes (Figures 7F and 7G). Proteins that were (de)phosphorylated in response to cholesterol in a RORα-dependent manner included S6 and LAMTOR1 (Figure 7H), aligning with our findings in the fly (Figures 5I and 5J), along with other key components of the TOR and insulin-signaling pathways including the TOR inhibitor TSC2, the complex KICKSTOR required for proper Rag modulation by nutrients, SIK3, the direct TOR kinase target 4EBP1, and the insulin/PI3K/PIP3 mediator PKB/AKT. These *in-vitro* findings also reinforce our *in-vivo* observations, suggesting a cell-autonomous modulation of TOR activity by RORα/HR3.

In results similar to those seen with *HR3* knockdown in the fly, 61 proteins were aberrantly responsive in terms of phosphorylation after 1 hour when RORα was silenced, indicating that RORα normally restrains these responses (Figures 7D and 7E). Following 6 hours of cholesterol stimulation, a larger array of proteins (318) showed cholesterol-induced phosphorylation changes only in RORα-deficient cells. This indicates the necessity of RORα/HR3 in maintaining homeostatic cellular responses to cholesterol over time. Collectively, our findings suggest that RORα and HR3 are essential for translating cholesterol abundance into TOR-pathway activation that governs cellular growth and metabolism (Figure 7I).

## Discussion

Organisms and cells must finely tune their growth in response to environmental fluctuations^11,40^. Since cholesterol is essential for cellular growth, the availability of cholesterol must be intimately linked with the activity of pathways that control growth. The TOR pathway is a primary growth-regulatory mechanism by which growth is adapted to nutritional cues. Our findings delineate dynamic regulation of the TOR pathway by varying cholesterol levels in *Drosophila* tissues, linking cholesterol availability with growth responses. This dynamic response requires a mechanism that accurately senses cholesterol levels and directly influences the activation of the TOR pathway. Our discovery suggests the nuclear receptor HR3 in *Drosophila* and its human ortholog RORα mediate this cholesterol-sensing step and translate sterol abundance into growth-regulating TOR activity. Cholesterol is a natural ligand of human RORα^41,42^, and our findings now show that cholesterol also acts as a *bona fide* ligand of its *Drosophila* ortholog, HR3. This represents a significant advance in understanding the sensing mechanism that links cholesterol availability to TOR and thus to cell growth.

Beyond showing that HR3 couples cholesterol levels to TOR regulation and to growth, our work also provides potential mechanistic links between steroid-regulated processes and nutrient-sensing pathways that coordinate growth during development^24,40,43,44^. In *Drosophila*, the interplay between the TOR/insulin and steroid (ecdysone) signaling pathways controls growth. Ecdysone antagonizes the growth of most larval tissues, but how it interacts with growth-regulatory pathways such as TOR has been an open question. *HR3* expression is induced by ecdysone^17^, and our work thus suggests a complex interplay in which ecdysone influences HR3 expression, which in turn adjusts growth according to cholesterol levels via TOR signaling. Furthermore, both HR3 and TOR regulate the biosynthesis of ecdysone^45–47^, and we recently provided evidence suggesting that cholesterol sensing in the PG is also linked to ecdysone production^11^. The dual function of HR3 in detecting levels of cholesterol – the precursor for ecdysone synthesis – and in stimulating TOR activity in the endocrine cells of the PG that produce ecdysone suggests that HR3 may directly link cholesterol sensing to steroid-hormone production, which governs both juvenile growth and the onset of maturation in animals.

HR3 function is also regulated by the nuclear receptor E75, which physically interacts with HR3 and represses its transcriptional activity. This interaction is regulated by the metabolic state of the PG – E75 is a heme-binding protein, and the oxidation state and gas binding of the iron ion govern E75:HR3 binding^48^. Interestingly, this regulatory nuclear-receptor pair not only operates in the fly but is conserved in humans, where their orthologs RORα and Rev-erb respond to the same respective ligands, cholesterol and heme/NO/redox^49–51^. Thus, these receptors may integrate different signaling cues into networks coordinating cellular growth processes in response to intracellular cholesterol levels. Our results corroborate earlier findings that HR3 genetically interacts with S6 kinase to modulate cell growth ^52^ and integrate cholesterol sensing into this growth-regulatory system. This regulation is mediated by an HR3 isoform lacking a DNA-binding domain, suggesting possible non-genomic effects. However, cholesterol likely binds to both isoforms of HR3, with and without the DBD. These isoforms likely have overlapping functions, and both could conceivably interact with E75 or other transcriptional regulators. Nuclear receptors often mediate effects via both transcriptional and non-genomic pathways, the latter involving direct interactions with signal-transduction pathway components at cytosolic or membrane locations^39^. We present evidence that cholesterol-induced TOR-pathway activation requires both transcriptional and non-transcriptional regulation and depends upon HR3. In humans, RORα regulates Wnt signaling through a non-genomic mechanism that involves Protein Kinase C α (PKCα) phosphorylation^53,54^. RORα is present in both the nucleus and cytosol, as well as in membrane domains rich in cholesterol. This suggests a potential for direct interaction of membrane-associated RORα with membrane cholesterol. An isoform of RORα lacking a DNA-binding domain has not yet been identified, leaving it an open question whether mammalian cells can produce an isoform similar to HR3. By acting via both transcriptional and non-transcriptional routes, HR3 and RORα may precisely regulate cholesterol-mediated growth responses. Intracellular signaling often involves counteracting feedback mechanisms to maintain homeostasis. Positive feedback amplifies signals, while negative feedback modulates these responses, preventing overactivation and allowing the cell to revert to its baseline state. Our results indicate that fat-tissue cells lacking HR3, when stimulated with cholesterol, develop high levels of intercellular variation in pS6, with some cells showing very high TOR pathway activity while neighboring cells exhibit low signaling. This suggests a loss of their ability to control the TOR pathway. An intriguing possibility is that HR3 modulates the TOR pathway through both positive and negative feedback mechanisms that involve rapid non-genomic responses and longer-term transcriptional regulation to maintain balanced cholesterol responses. Recent research has implicated LYCHOS and SLC38A9 in the activation of the TOR pathway by lysosomal cholesterol ^3,14^. These factors act upstream of the Rag GTPases but are not reported to be involved in any negative feedback regulation. Our findings indicate the additional necessity of HR3/RORα for precise modulation of TOR activity, critical not only for immediate responses to cholesterol but also for sustained pathway equilibrium. Thus, HR3/RORα may act as a gatekeeper, ensuring that while cholesterol effectively stimulates growth via TOR, it does not induce overactivation that could lead to pathological hyperactivation which could lead to tumorigenesis.

Our study may thus also shed light on the link between high cholesterol and cancer development and progression. Extensive studies have reported an association between high levels of cholesterol in the blood and an increased risk of various cancers^7,55,56^. Although the exact molecular mechanisms connecting elevated cholesterol to cancerous cell proliferation remain elusive, it is known that the enhanced uptake of exogenous cholesterol by cells stimulates oncogenic processes and tumor growth^10^. Our findings show that in response to cellular uptake of exogenous cholesterol, RORα regulates numerous signaling pathways involved in cancer initiation and progression, such as IGF, PI3K/AKT, MAPK, ALK, and SIK3. SIK3 expression is highly elevated in the majority of breast cancer cases^57^, which are typically characterized by their dependence on cholesterol^58^. This suggests that RORα and TOR signaling may mechanistically link cholesterol and tumor growth.

## Methods

### *Drosophila* media and husbandry

Flies were kept at 25 °C and 60% humidity with a 12-hour/12-hour daily light cycle. Fly stocks were maintained on standard fly food (STD-FF) medium containing 6% sucrose, 0.8% agar, 3.4% yeast, 8.2% cornmeal, 0.16% Tegosept antifungal agent, and 0.48% propionic acid preservative. Unless a different diet is indicated, this medium was also used for experiments. Fly lines used are described in the Key Resources Table.

### Key Resources Table

| REAGENT or RESOURCE | SOURCE | IDENTIFIER |
| --- | --- | --- |
| **Antibodies** |  |  |
| Anti-GFP, mouse clone 3E6 | ThermoFisher | A11120<br>RRID AB_221568 |
| Anti-guinea pig IgG, Alexa Fluor 488 conjugate | ThermoFisher | A21450<br>RRID AB_2735091 |
| Anti-histone H3, rabbit | Abcam | 1791; No RRID |
| Anti-HR3, rabbit | Gift from J. Montagne<br>(Friedrich Miescher Institute<br>for Biomedical Research).<br>(Montagne J *et al.*<sup>32</sup>) | n/a |
| Anti-mouse IgG, Alexa Fluor 488 conjugate | ThermoFisher | A32723<br>RRID AB_2633275 |
| Anti-phospho-S6 (*Drosophila*) | Romero-Pozuelo J *et al.*<sup>27</sup>;<br>Texada MJ *et al.*<sup>13</sup> | n/a |
| Anti-phospho-S6 (Human) | Cell Signaling | 4857 |
| Anti-phospho-S6K (*Drosophila*, pT398), rabbit | Cell Signaling | 9209S |
| Anti-rabbit IgG, Alexa Fluor 488 conjugate | ThermoFisher | A32731<br>RRID AB_2633280 |
| Anti-rabbit IgG, Alexa Fluor 555 conjugate | ThermoFisher | A32732<br>RRID AB_2633281 |
| Anti-rabbit IgG, Alexa Fluor 647 conjugate | ThermoFisher | #A32733<br>RRID AB_2633282 |
| Anti-RORα (human) Rabbit mAb (E6G51) | Cell Signaling | 34639S |
| Anti-S6 – western blot | Cell Signaling | 2317 |
| Anti-Shroud, guinea pig | Gift from R. Niwa (University<br>of Tsukuba). (Shimada-Niwa Y<br>and Niwa R<sup>59</sup>) | n/a |
| Anti-α-Tubulin, mouse | Developmental Studies<br>Hybridoma Bank | AA4.3 |
| Goat anti-mouse IgG (H+L) peroxidase<br>conjugate | Jackson ImmunoResearch | 115-035-003 |
| Goat anti-rabbit IgG (H+L) peroxidase<br>conjugate | Jackson ImmunoResearch | 111-035-003 |
| Goat anti-rabbit IgG (H+L) peroxidase<br>Conjugate | ThermoFisher | 31466 |
| IRDye 680RD-labeled goat anti-mouse | Li-COR | 925-68070 |
| IRDye 800CW goat anti-rabbit | Li-COR | 925-32210 |
| **Chemicals, peptides, and recombinant proteins** |  |  |
| 20-hydroxyecdysone (20E, ecdysone) | Sigma-Aldrich | H5142 |
| 2x Laemmli sample buffer | Bio-Rad | 1610737 |
| Agar, bacteriological grade | VWR | J637-1KG |
| Benzonase | Sigma-Aldrich | G9422 |
| β-Glycerophosphate disodium salt | Sigma-Aldrich | 154808-51-0 |
| Bicinchoninic acid solution | Sigma-Aldrich | B9643-1L |
| Biotin | Sigma-Aldrich | B4501; CAS 58-85-5 |
| Bovine serum albumin (protease-free, lyophilized) | BioWest | P6155-1KG |
| Casein | Sigma-Aldrich | C5679; CAS 9000-71-9 |
| Chameleon Duo pre-stained protein ladder | Li-COR | 928-60000 |
| DAPI | ThermoFisher | 62248; CAS 28718-90-3 |
| Chloroform | Carlo Erba Reagents | 438612 |
| Cholesterol | Sigma-Aldrich | C3045; CAS 57-88-5 |
| Cholesterol | Steraloids | C6807-000 |
| Cholesterol sulfate | Steraloids |  |
| Choline chloride | Sigma-Aldrich | C1879; CAS 67-48-1 |
| Copper(II) sulfate solution | Sigma-Aldrich | C2284-25ML |
| Cyloheximide | Sigma-Aldrich | 239765 |
| 7-dehydrocholesterol | Steraloids | C3000-000 |
| DNase | Macherey-Nagel | 740955.250 |
| D-Pantothenic acid, hemicalcium salt | Sigma-Aldrich | P5155; CAS 137-08-6 |
| Farnesol | Sigma | 93547 |
| Farnesyl acetate | Sigma | 340480 |
| Fetal bovine serum (FBS, Qualified, HI) | Gibco | 10500-064 |
| Fetal bovine serum | Wisent | 090-110 |
| Folic acid | Sigma-Aldrich | F8758; CAS 59-30-3 |
| Geraniol | Sigma | 163333 |
| GSK157369x | GlaxoSmithKline |  |
| Imidazole | Sigma | I2399 |
| Inosine | Sigma-Aldrich | I4125; CAS 58-63-9 |
| Intercept blocking buffer | Li-COR | 927-70001 |
| IPTG | Bioshop | IPT002 |
| Lipofectamine 2000 | Invitrogen | 11668-030 |
| Lipoprotein deficient serum from fetal calf (Lipid-depleted serum, LDS) | Sigma-Aldrich | S5394-50ML |
| Methoprene | Sigma | 33375 |
| Methyl-β-Cyclodextrin (MCD) | Sigma-Aldrich | C4555 |
| M3 medium (Shields and Sang) | Sigma | S3652 |
| Nicotinic acid | Sigma-Aldrich | N0761; CAS 59-67-6 |
| NutriFly medium | Genesee Scientific | "Bloomington formula," 66-112 |
| N,O-BSTFA | Sigma | 155195 |
| Opti-MEM (1X) reduced-serum medium | Gibco | 31985-062 |
| Penicillin-Streptomycin (Pen-Strep) | Gibco | 15140-122 |
| Phalloidin, Alexa Fluor 488 conjugate | ThermoFisher | A12379 |
| Phalloidin, Alexa Fluor 647 conjugate | ThermoFisher | A22287 |
| PhosStop – phosphatase inhibitor tablets | Roche | PHOSS-RO |
| PMSF | Sigma | 10837091001 |
| Poly-lysine | Sigma-Aldrich | P8920-100ML |
| ProLong Glass Antifade Mountant | Life Technologies | P36984 |
| Propanediol | Sigma | 12279 |
| Pyridoxine | Sigma-Aldrich | P6280; CAS 58-56-0 |
| Riboflavin | Sigma-Aldrich | R4500; CAS 83-88-5 |
| Roche Complete Protease Inhibitor cocktail with EDTA (tablets) | Roche | 11836153001 |
| Schneider's *Drosophila* Medium (1x) | Gibco | 21720 |
| Skim milk powder | Millipore | 70166-500G |
| Squalene | Sigma | S3626 |
| StableCell RPMI-1640 | Sigma-Aldrich | R2405-500ML |
| Stearic acid | Sigma | 175366 |
| Superdex 16/60 resin | GE Healthcare | n/a |
| TCEP | Sigma | C4706 |
| Tegosept (Methyl 4-hydroxybenzoate) | VWR | 2505.293 |
| Thiamine | Sigma-Aldrich | T1270; CAS 67-03-8 |
| Trypsin-EDTA (0.25%) | Gibco | 25200056 |
| Tween20 | Sigma-Aldrich | P9416-100ML |
| Uridine | Sigma-Aldrich | U3003; CAS 58-96-8 |
| VectaShield Antifade Mounting Medium with DAPI | Vector Laboratories | H-1200 |
| **Critical commercial assays and kits** |  |  |
| RNeasy Mini Kit | Qiagen | 74104 |
| Steady-Glo Luciferase Assay System | Promega | E2510 |
| Glo Lysis Buffer | Promega | E2661 |
| Trans-Blot Turbo Midi 0.2 µm Nitrocellulose Transfer Packs | Bio-Rad | 1704159 |
| ECL Prime Western Blotting Detection Reagents | Amersham | RPN2232 |
| 10% Mini-PROTEAN TGX precast polyacrylamide gel | Bio-Rad | 4561034 |
| 12% Mini-PROTEAN TGX precast polyacrylamide gel | Bio-Rad | 4561044 |
| 4-20% Mini-PROTEAN TGX Precast polyacrylamide gel | Bio-Rad | 4561094 |
| SecureSeal imaging spacers | Grace Bio-labs | GBL654006-100EA |
| **Experimental models: cell lines** |  |  |
| Karpas-707H cells | University of Cambridge (Karpas A *et al.*<sup>60</sup>) | https://www.enterprise.cam.ac.uk/reagents/karpas-707h-human-myeloma-cell-line/ |
| Hi5 insect cells | ThermoFisher | #B85502 |
| **Experimental models: *Drosophila* stocks** |  |  |
| *Cg-GAL4* | Bloomington *Drosophila* Stock Center (BDSC) (Asha H *et al.*<sup>61</sup>) | 7011 |
| *Heat-shock-GAL4(DBD)::Hr3(LBD);UAS-GFP* | BDSC (Palanker L *et al.*<sup>34</sup>) | 28868 |
| *phm-GAL4 TubGAL80<sup>TS</sup>* | n/a | Made from BDSC 7018 |
| *phm-GAL4; Tub-GAL80<sup>TS</sup>* | McGuire SE *et al.*<sup>62</sup> | Made from BDSC 7108 and *phm*<sup>#22</sup>-GAL4 (Gift from M. O'Connor, University of Minnesota) |
| *pumpless (ppl)-GAL4* | BDSC (Zinke I *et al.*<sup>63</sup>) | 58768 |
| *Tub-GAL80<sup>TS</sup>; da-GAL4* | n/a | Made from BDSC 7108 and 55850 |
| *Tub-GAL80<sup>TS</sup>; Tub-GAL4* | n/a | 86328 |
| *UAS-EcR-RNAi* | BDSC (Ni *et al.*<sup>64</sup>) | 37058 and 37059 |
| *UAS-Hr3(LBD)* | Gift from J. Montagne. (Montagne J *et al.*<sup>32</sup>) | DHR3-RS |
| *UAS-Hr3-RNAi* | Vienna *Drosophila* Resource Center (VDRC). (Dietzl G *et al.*<sup>65</sup>) | 10687 |
| *UAS-Hr3-RNAi<sup>GD</sup>* | VDRC | 12044 |
| *UAS-Hr3-RNAi<sup>TRIP</sup>* | BDSC | 27253 |
| *UAS-Npc1a-RNAi<sup>KK</sup>* | VDRC | 105405 |
| *UAS-RagA-B-RNAi* | BDSC | 34590 |
| *UAS-RagC-D-RNAi* | BDSC | 32342 |
| *UAS-S6K-RNAi* | VDRC | 104369 |
| *UAS-S6K<sup>STED</sup>* | BDSC | 6913 |
| *UAS-Tor-RNAi* | BDSC | 34639 |
| *unk-FireflyLuciferase (B76)* | Gift from A. Teleman (Tiebe M *et al.*<sup>30</sup>) | n/a |
| *w<sup>1118</sup>* | VDRC | 60000 |

| Oligonucleotides |  |  |
| --- | --- | --- |
| Human RORα siGenome SMARTpool | Horizon - Dharmacon | M-003440-01-0005 |
| Software and algorithms |  |  |
| CellProfiler 3.1.9 | http://cellprofiler.org/ | McQuin *et al.*<sup>66</sup> |
| FIJI Image-processing package | http://fiji.sc/ | Schneider *et al.*<sup>67</sup> |
| PANTHER – Classification system – version 18.0 | pantherdb.org | Thomas *et al.*<sup>68</sup> |
| Prism 10 | GraphPad | n/a |
| SRplot | GO, Pathway enrichment<br>bubblor plot<br>[SRplot - Free online GO Pathway bubble plot (bioinformatics.com.cn)](http://SRplot-FreeonlineGOPathwaybubbleplot%28bioinformatics.com.cn%29) | Tang *et al.*<sup>69</sup> |
| STRING: functional protein association networks ; version 12.0 | [STRING: functional protein association networks (string-db.org)](http://STRING:functionalproteinassociationnetworks%28string-db.org%29) | Szklarczyk *et al.*<sup>70</sup> |
| ZEN 3.5 (ZEN lite) | Zeiss | n/a |
| Xcalibur v3.0 | ThermoFisher Scientific | n/a |
| MS Workstation | Varian | n/a |
| Mascot v 2.2.04 | Matrix Science Ltd. | n/a |
| Proteome Discoverer v2.5.0.400 | ThermoFisher Scientific | n/a |
| Mass Lynx 4.0 | Micromass | n/a |
| Other |  |  |
| AKTA FPLC apparatus | GE Healthcare | n/a |
| 3900 gas chromatograph | Varian | n/a |
| Saturn 2100T ion-trap mass spectrometer | Varian | n/a |
| Quadrupole time-of-flight mass spectrometer | Micromass | n/a |
| Fused silica column | Varian | CP8944 |
| Tandem mass tags 18-plex | ThermoFisher Scientific | A52045 |
| EASY nanoLC | ThermoFisher Scientific | n/a |
| Fusion Lumos Tribrid mass spectrometer | ThermoFisher Scientific | n/a |
| Orbitrap Eclipse Tribrid mass spectrometer | ThermoFisher Scientific | n/a |

### Timed egg-lays

Virgin females from the driver stocks were mated with males of the RNAi lines in bottles containing STD-FF and additional dry-yeast pellets approximately 2-days prior to timed egg-lays. The day before egg collection began, animals were transferred to egg-laying chambers with apple-juice/agar egg-collection plates supplemented with yeast paste. Egg-collection plates were changed on the following day for egg collection during a 4-hour period (ZT1 to ZT5) at 25 °C at 60% humidity. For genotypes not carrying *GAL80^TS^*, first-instar hatchers were collected after 24 hours at 25 °C and transferred to vials containing STD-FF or NutriFly (NF) at 30 larvae per vial containing 10 mL of medium. Offspring carrying *Tub-GAL80^TS^* were collected after a 48-hour hatching period at 18 °C to prevent GAL4 activity during early larval development.

### Chronic high-cholesterol diet

Bloomington NutriFly (NF) medium (Genesee Scientific, “Bloomington formula”, #66-113), which contains a relatively low yeast/sterol content, was cooked according to product guidelines with added 0.16% Tegosept antifungal agent. The medium was supplemented while liquid with cholesterol (from a 8-mg/mL stock of cholesterol in ethanol) to a final added concentration of 40 μg/mL (NF+40) or 80 μg/mL (NF+80). Control media were prepared by adding the same amount of 100% ethanol. First-instar larvae were picked from egg collections and developed in these media until experiments were conducted during the third instar.

### Synthetic-food re-feeding experiments: Cholesterol, protein, and ecdysone

The synthetic food (SF) diet is described in Reis, 2016 ^23^, adjusted to increase its nutritional content and solidity. Casein was used at double the concentration, agar at 1% concentration, sucrose was increased to 2.7-fold, and choline chloride was doubled. The final components of 1 L of food are 10 g lipid-depleted agar, 146.6 g lipid-depleted casein, 35.5 g sucrose, 0.64 g choline chloride, 0.85 g inosine, 0.76 g uridine, 133 mL of 10 g/L NaHCO_3_ stock, 133 mL of a 37.3 g/L KH_2_PO_4_ stock), 133 mL of 7.1 g/L K_2_HPO_4_ stock, 133 mL of 6.2-g/L MgSO_4_♦7H_2_O stock, 133 mL of Vitamin A stock, 13.3 mL of Vitamin B stock, 333 mL deionized water, 13.3 mL of 10% Tegosept in ethanol. Vitamin A stock contained 20 mg thiamine, 100 mg riboflavin, 120 mg nicotinic acid, 160 mg D-pantothenic acid hemicalcium salt, 25 mg pyridoxine, 2 mg biotin in 1 L deionized water. Vitamin B stock was made of 500 mg folic acid dissolved in 167 mL 20% ethanol. Cholesterol content was adjusted by adding either pure ethanol or 8 mg/mL cholesterol in ethanol to reach final cholesterol concentrations of 0, 1.2, 6, 15, 25, 40, or 80 µg/mL. Casein and agar were lipid-depleted through six chloroform extractions of at least 6 hours each, with stirring, at room temperature, at a 1:5 ratio of solids to chloroform. After each extraction, solid material was collected by filtration (using Whatman paper in a ceramic funnel), and lipid-free material was dried after the last collection by allowing chloroform to evaporate under a fume hood.

Larvae from timed egg collections were developed on STD-FF till early third instar: genotypes without GAL80^TS^, 25 °C until 84 hours AEL; those with GAL80^TS^, 18 °C until 120 h AEL. Larvae were then collected by floating in 20% sucrose solution, washed using DI water, and transferred using a paint brush or entomological forceps to synthetic media in a 24-well dish, each well stoppered with a foam plug. For cholesterol- and ecdysone-feeding experiments, animals were transferred at this time to media containing either 0 µg/mL or 1.2 µg/mL cholesterol (with other ingredients constant); for protein-feeding experiments, this medium lacked casein but did contain 80 µg/mL cholesterol. After 10-12 hours on this medium, the larvae were collected using DI water and transferred either to fresh nutrient-dropout medium (0 or 1.2 µg/mL cholesterol or 0 mg/mL protein) or to a re-feeding medium as indicated in each figure (containing 40 or 80 µg/mL cholesterol, 250 µg/mL 20-hydroxyecdysone + 1.2 µg/mL cholesterol, or 14.7 mg/mL casein as appropriate) in new 24-well plates. After the indicated time intervals (15 minutes, 30 minutes, or 1, 2, 4, 6, or 10 hours), the larvae were collected using water washes.

### Larval-mass measurements

First-instar larvae were collected from timed egg-lays, transferred to the indicated diets, and maintained at 25 °C. At 96 h AEL, larvae were collected and rinsed with deionized water. Batches of larvae were chilled in DI water on ice, and individual animals were blotted dry on a KimWipe and weighed using a Sartorius SE2 Micro Balance.

### Promega Steady-Glo Luciferase assay

Three feeding late-third-instar larvae for each sample were collected into Glo Lysis Buffer (Promega, E2661) in 2-mL Eppendorf tubes. For normal assays, 200 µL of buffer was used per sample; for large animals or if protein quantification was performed in parallel, 400 µL was used. The samples were homogenized with a 5-mm steel bead in a TissueLyser bead mill (Qiagen; operated at 50 Hz for 30 seconds). The homogenized samples were incubated at room temperature for 10 minutes for complete cell lysis and then centrifuged at maximum speed for 5 minutes to pellet insoluble material. An aliquot (150 µL) of supernatant was removed to new Eppendorf tubes for protein quantification and placed on ice. For luciferase measurement, 20 µL of each sample was transferred at room temperature to an opaque white 96-well plate (Costar), and 20 µL of Steady-Glo Luciferase Reagent (Promega, E2510) was added to each well, with gentle mixing by pipetting. The plate was centrifuged to settle the liquids and incubated at room temperature in the dark for 10 minutes to allow the luciferase reaction to reach a steady state, after which the luminescence was measured using an EnSight multi-mode plate reader (PerkinElmer).

### Protein Quantification (Bicinchoninic Acid Method)

The protein concentrations in the aliquots set aside during the luciferase assay and for human western-blot protein-loading calculations were measured using a bicinchoninic acid (BCA)-based method. BCA working reagent was prepared by combining bicinchoninic acid solution (Sigma-Aldrich, #B9643) and 4% cupric sulfate solution (Sigma-Aldrich, #C2284) at a 50:1 ratio. Sample supernatant was diluted by adding of 3 volumes of PBS, and four microliters of the diluted material were pipetted into a 384-well plate. Thirty microliters of BCA working reagent was added to each well, with gentle mixing by pipette, and the plate was centrifuged to settle liquids and incubated for 30 minutes at 37 °C. The absorbance of each sample at 540 nm was measured using an EnSight multi-mode plate reader (PerkinElmer).

### Western blotting of *Drosophila* samples

Three to eight feeding (pre-wandering) late-third-instar larvae for each sample were lysed in ice-cold SDS sample buffer (Bio-Rad, 2x Laemmli Sample Buffer, #1610737), containing 5% beta-mercaptoethanol and protease and phosphatase inhibitors (Roche Complete Mini protease inhibitor, Sigma-Aldrich #11836153001, and Roche Complete Ultra phosphatase inhibitor, Sigma-Aldrich #05892970001), 60 µL per larva, using a TissueLyser bead mill (Qiagen). Samples were denatured at 95 °C for 5 minutes, and insoluble material was pelleted by centrifugation at top speed for 5 minutes. Samples were loaded into precast 4%-20% gradient polyacrylamide gels (Bio-Rad, #4561094) and electrophoresed at 150 V for approximately 35 minutes. Proteins were transferred to 0.2-µm nitrocellulose membrane using a Trans-Blot Turbo Transfer Pack (Bio-Rad, #1704159) and the Bio-Rad dry-transfer apparatus. Membranes were then blocked in Intercept Blocking Buffer (LI-COR, #927-70001) for 1 hour at room temperature with gentle agitation. Phospho-S6K (pS6K) and histone H3 were detected by incubating with rabbit anti-pS6K (Cell Signaling #9209S, diluted 1:1000) and rabbit anti-histone-H3 (Abcam #1791, diluted 1:1000) in Odyssey blocking buffer (LI-COR) + 0.2% Tween-20 (Sigma-Aldrich, #P9416) overnight at 4 °C with gentle agitation. Membranes were washed three times with PBS+0.1% Tween-20, and secondary staining was performed with IRDye 680RD-labeled goat anti-mouse and IRDye 800CW conjugated goat anti-rabbit (LI-COR, #925-68070 and #925-32210, each diluted 1:10,000 in Intercept Blocking Buffer + 0.2% Tween-20) for 45 minutes at room temperature with gentle agitation. Membranes were washed three times with PBS+0.1% Tween-20 followed by a single wash with PBS. Bands were visualized using an Odyssey Fc gel reader (LI-COR) and quantified using the LI-COR Image Studio Gel Reader program.

### Immunostaining, microscopy, and quantification

Larval tissue was dissected in PBS and fixed in fresh 4% paraformaldehyde (EM grade) in PBS at room temperature: for fat-body tissue, 10-15 larvae were inverted and fixed at room temperature for 45 minutes, whereas for prothoracic-gland (PG) samples, tissues were accumulated in ice-cold 4% PFA during dissection and fixed at room temperature for 70 minutes. Samples were quickly rinsed in PBST (PBS + 0.1% Triton X-100) and washed three times for 15 minutes in PBST. The samples were then blocked in PBST+3% normal goat serum (Sigma) for at least 30 minutes at room temperature with gentle agitation. Tissues were incubated with primary antibodies diluted in PBST + 3% normal goat serum overnight at 4 °C with gentle agitation. Tissue was then washed three times in PBST. Samples were incubated with secondary antibodies diluted in PBST at 4 °C, in the dark, overnight with gentle agitation. For actin staining, samples were incubated with phalloidin (Alexa Fluor 647 conjugate, ThermoFisher #A22287, or Alexa Fluor 488 conjugate, #A12379, diluted 1:100 in PBST for 1 hour), and for nuclear staining, samples were incubated in DAPI (1:500 in PBS, ThermoFisher, #62248) for 0.5-1 hour. Samples were washed twice in PBST and once in PBS and mounted on glass slides coated twice with poly-L-lysine (Sigma-Aldrich, #P8920-100ML). The mounted tissue was imaged using a Zeiss LSM 900 confocal microscope using a 20x objective (NA 0.8). All samples compared within a figure panel were processed similarly (fixation, staining preparations and imaging hardware settings). In all cases, each experimental sample was prepared simultaneously with its respective control using the same reagent preparations. Data sets from multiple preparations are normalized to controls before comparison.

Primary antibodies used were rabbit anti-pS6^27^ against the epitope RRR(phospho-S)A(phospho-S)IRE(phospho-S)K, used at 1:500; mouse anti-GFP (clone 3E6, ThermoFisher #A11120, RRID AB:221568, at 1:500); guinea-pig anti-Shroud^59^ (a kind gift from R. Niwa, University of Tsukuba; 1:200); and rabbit anti-HR3^32^, a generous gift from J. Montagne (Friedrich Miescher Institute for Biomedical Research, Switzerland, 1:250). Secondary antibodies (ThermoFisher, 1:500) were all raised in goats and were cross-adsorbed by the manufacturer to reduce off-target binding. These included anti-rabbit, Alexa Fluor 488 conjugate (#A32731, RRID AB_2633280); anti-rabbit, Alexa Fluor 555 conjugate (#A32732, RRID AB_2633281); anti-rabbit, Alexa Fluor 647 conjugate (#A32733, RRID AB_2633282); anti-guinea-pig, Alexa Fluor 647 conjugate (#A21450, RRID AB_2735091); and anti-mouse, Alexa Fluor 488 conjugate (#A32723, RRID AB_2633275).

For quantification of PG cell size and pS6 intensity, a composite of channels was created, and individual cells and their nuclei (based on actin and DAPI staining) were traced at the center of the cell thickness in the Z-stack, saving this to the region-of-interest (ROI) manager in the image-analysis package FIJI ^67^. The intensity of staining in each channel was quantified in each ROI. The pS6 intensity in the cytoplasm was calculated as

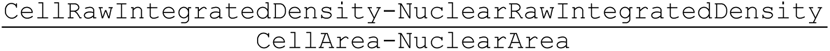

For quantification of fat-body pS6, a large number of tissues were mounted on each slide. Using a 5x overview scan in the DAPI channel on the Zeiss LSM 900 confocal microscope, areas of flat, single-cell-thick tissue were selected for each large piece of fat body and were imaged with the 20x objective with a 1.5-μm Z spacing. Z-stacks were pre-processed in FIJI using a macro for creating a composite image and Z-projected using “sum”. Areas of folded or overlaid tissues were removed by manual segmentation; the channels were split, and a binary mask was created from the DAPI channel. This mask was used to remove the nucleus from the pS6 channel, since nuclear pS6 staining is believed to be spurious^27^. Quantification was automated using CellProfiler, version 3.1.9^66^: each cell’s pS6 signal was quantified by segmentation of the tissue into individual cells using the actin (phalloidin staining) channel and the binary mask created using DAPI in order to produce a cytoplasmic pS6 signal. The pS6 signal was measured as the average intensity across each area (“integrated intensity” divided by the cytoplasmic area).

### *Ex-vivo* culture of *Drosophil*a fat body

Larvae from a four-hour egg collection were raised on STD-FF at 25 °C. At 96 h AEL, fat-body tissue was dissected in Schneider’s insect medium (Gibco, #21720). Tissue was cholesterol-depleted using Schneider’s medium containing 10% lipid-depleted-serum (LDS, Sigma-Aldrich, #S5394-50mL) and 0.75% methyl-β-cyclodextrin (MCD, Sigma-Aldrich, #C4555), for 2 hours at room temperature with gentle rocking in a glass staining dish with glass cover (Assistant, #42020010). For experiments including cycloheximide, 15 minutes prior to cholesterol stimulation, cholesterol-depletion medium was replaced with fresh cholesterol-depletion medium containing 100 μg/mL cycloheximide (Sigma-Aldrich, #239765) from a 100-mg/mL stock solution in DMSO. Tissue was washed twice with room temperature Schneider’s medium and then incubated in cholesterol-depletion or cholesterol-stimulation medium for 1 hour. Conditions including cycloheximide also contained it during the stimulation period. Cholesterol-stimulation medium consisted of Schneider’s medium containing 10% LDS, 0.2% MCD, and 100 µM (39 µg/mL) cholesterol from a 20-mg/mL cholesterol stock in ethanol. After incubations, media were replaced by 4% PFA for 45-minute tissue fixation at room temperature. Fixed tissues were then processed for immunostaining as usual.

### Human cell culture

Karpas 707h cells^60^ (Cambridge University; Cambridge Enterprise Limited) were cultured in StableCell RPMI-1640 culture medium (Sigma-Aldrich, #R2405-500ML) containing 15% fetal bovine serum (Gibco, #10500-064) and Pen-Strep antibiotic mixture (100 u/mL Penicillin, 100 µg/mL Streptomycin, Gibco, #15140-122). The cultures were maintained at 37 °C and 5% CO_2_. Karpas cells partly attach to plasticware after 3-4 days of culturing, while some cells remain in suspension. To prevent selective pressure during passaging (cells can be partly released with tapping on cultureware), Karpas cells were passaged by collecting cells in suspension and using Trypsin-EDTA (Gibco, #25200056) to fully float attached cells.

### Cell-culture cholesterol starvation/re-stimulation

The methods used are similar to those described in Shin et al.^37^ and Castellano et al.^3^ Cells were cholesterol-depleted using cell-culture media supplemented with methyl-β-cyclodextrin (MCD, Sigma-Aldrich, #C4555) and lipid-depleted serum (Lipoprotein Deficient Serum from fetal calf, LDS, Sigma-Aldrich, #S5394-50mL). Cells were re-stimulated with cholesterol using cell-culture medium supplemented with MCD:cholesterol complex and LDS.

Cells were plated in standard growth medium, RPMI-1640 medium containing 15% fetal bovine serum and Pen-Strep, in 6-well plates (Greiner bio-one, #657160) and allowed to attach for 3-4 days. Cells were then rinsed with serum-free RPMI-1640 medium and then incubated with 0.75% MCD with 0.5% LDS in RMPI-1640 (without antibiotics) for 2 hours in order to cholesterol-deplete the cells. Cells were re-stimulated with cholesterol in RPMI-1640 using complexed MCD:cholesterol (0.1% MCD and 50 µM/∼20 µg/mL cholesterol or a 2x dose of 0.2% MCD and 100 µM cholesterol) plus 0.5% LDS. MCD:cholesterol complex was prepared by diluting a 20-mg/mL cholesterol stock in ethanol into a 15-mL Falcon tube containing RMPI-1640, 0.1% MCD, and 0.5% LDS, resulting in a 50-uM final concentration of both cholesterol and MCD. The tube was then vortexed and incubated in a 37 °C water bath for 2 hours before being applied to the cells.

### Knockdown of *RORα* using siRNA

Knockdown of *RORα* in human cells was performed using the siGenome SMARTpool Human RORA (#6095) siRNA from the Horizon Discovery “Dharmacon” portfolio (#M-003440-01-005) and Lipofectamine 2000 (Invitrogen, #11668-030). The siRNA stock was diluted to 20 µM in RNase-free water following the product protocols. A siRNA duplex targeting *Renilla Luciferase* (a gene not present in these cells) was used as a control. siRNA and Lipofectamine solutions were prepared according to product specifications. Cells were rinsed with serum-free RPMI-1640 medium before being transfected with 5 µL of siRNA pool (20 µM) and 5 µL Lipofectamine 2000 in a 6-well plate in 1 mL of antibiotic-free Opti-MEM (Gibco, #31985-062) medium plus 2 mL antibiotic-free cell-culture medium. For experiments other than phosphoproteomics, the medium was changed 24 hours after treatment to new RPMI-1640 with 15% fetal bovine serum (without antibiotics). Seventy-two hours after knock-down treatment, cells were processed for further experiments (cholesterol treatments or collection of samples). Samples for phosphoproteomics were exposed to the siRNA treatment for the entire 72 h incubation, with additional siRNA and culture medium being added after 24 h for a stronger knock-down.

### Human-cell lysis and immunoblotting

Cells in suspension were collected using RPMI-1640 and centrifuged for 4 minutes at 900 rpm, and the supernatant was removed. Lysis buffer was added to still-attached cells in the culture flask and to the pelleted suspension quickly after medium was removed. Cells were lysed in ice-cold RIPA buffer (50 mM Tris-HCl pH 7.5, 150 mM sodium chloride, 0.5% sodium deoxycholate, 1% Nonidet P-40, 0.1% SDS) supplemented with Roche Complete Protease Inhibitor Cocktail with EDTA (Roche, #11836153001), PhosStop (used at 2x, Roche, #PHOSS-RO), 11 mg/mL β-glycerophosphate (from a 110-mg/mL 10x stock solution), ∼100 mM sodium fluoride (a saturated solution in RIPA buffer, circa 1 M, was made as a 10x stock solution), and Benzonase endonuclease (Sigma-Aldrich, #G9422). Cells were transferred to pre-cooled Eppendorf tubes on ice. The samples were incubated on ice for 10 minutes after which they were spun down to pellet nuclei and membrane at maximum speed (14,000 rpm) at 4 °C for 15 minutes. An aliquot of supernatant was reserved for BCA protein measurement. For Western samples, 230 μL of supernatant was mixed with 70 μL 5x Laemmli buffer (4% SDS, 10% β-mercaptoethanol, 20% glycerol, 0.004% bromophenol blue, 0.125 M Tris-HCl, pH 6.8) or 230 μL 2x Laemmli Sample Buffer (Bio-Rad, #1610737), and samples were immediately denatured for 5 minutes at 95 °C.

Loaded sample sizes were adjusted to equalize total protein based on BCA protein measurements. Protein separation was done using electrophoresis in polyacrylamide gels using 1x Running Buffer (10x running buffer: Tris base 30.2 g, glycine 188 g, 10% SDS solution 100 mL, in deionized water to 1 L) using a BioRad Mini-PROTEAN Tetra Vertical Electrophoresis Cell system. Protein mass ladder PageRuler Plus Prestained Ladder (Thermo Scientific, #26615) or Chameleon Duo Pre-stained Protein Ladder (LI-COR, #928-60000) was used. The gel in Figure 7B was run on a 12% Mini-Protean TGX precast protein gel (Bio-Rad, #4561034), whereas the one in Figure 7C was run using hand-cast gels. Gels were run at 20 mA per gel at max voltage (300 V) for approximately 1 hour. The blot in Figure 7B was transferred using semi-dry transfer as described in the *Drosophila* western-blot section, whereas Figure 7C was transferred to 0.2-µm nitrocellulose membrane (Amersham Proteon Nitrocellulose blotting membrane) using wet transfer: the blotting was performed using 1x Running Buffer with 20% methanol, kept cold with ice packs, run at 100 V for 1 hour. Ponceau stain was used to visualize the transferred proteins.

Membranes were blocked for 1 hour at room temperature using PBS + 5% nonfat milk powder + 0.1% Tween-20 and stained overnight at 4 °C in primary-antibody mix (diluted in PBS + 0.1% Tween- 20 + 5% bovine serum albumin). Rabbit anti-pS6 (Cell Signaling, #4857) was used at 1:1000, mouse anti-S6 (Cell Signaling, #2317) was used at 1:1000, and mouse anti-α-Tubulin (University of Iowa Developmental Studies Hybridoma Bank clone #AA4.3) was diluted 1:5000. Membranes were washed three times for 15 minutes in PBS+0.1% Tween-20 at room temperature before secondary staining. Secondaries (goat anti-rabbit IgG H+L, HRP conjugate (ThermoFisher, #31466, or Jackson ImmunoResearch, #111-035-003) and goat anti-mouse IgG H+L, HRP conjugate (Jackson ImmunoResearch, #115-035-003) were diluted 1:10,000 in PBS + +5% nonfat milk powder + 0.1% Tween-20 at 1:10,000 and incubated on the blot for 1-2 hours at room temperature. Stain was detected using SuperSignal West Femto Maximum Sensitivity Substrate (Thermo Scientific, #34094) or ECL Prime Western Blotting Detection Reagents (Amersham, #RPN2232) chemiluminescence using a Bio-Rad ChemiDoc Touch Imaging System or an Odyssey Fc gel reader (LI-COR).

### RNA sequencing

*Tor* or *HR3* were knocked down in *Drosophila* larvae using the temperature-inducible ubiquitous driver *Tub-GAL80^TS^; Tub-GAL4*. An 8-hour egg collection was made at 18 °C to prevent activation of the GAL4 system, and egg-laying plates were maintained at 18 °C for an additional 44 hours before first-instar larvae were collected using a metal probe into vials containing standard food (30 larvae per vial). The collected larvae were maintained at 18 °C for a further 72 hours before being transferred to 29 °C to induce RNAi expression (at 120 hours AEL). After nine hours of RNAi induction on standard food (at 129 hours AEL), larvae were transferred to low-cholesterol (1.2 µg/mL) synthetic medium and incubated at 29 °C for 11 hours more (until 140 hours AEL). Larvae were then transferred to synthetic food containing either 80 µg/mL cholesterol (for replenishment) or 1.2 µg/mL (control) and incubated at 29 °C for 10 hours. At this time (150 hours AEL), five feeding late-third-instar larvae were collected for each of five replicates into ice-cold RLT buffer (Qiagen RNeasy Mini kit, #74104) containing 1% β-mercaptoethanol and homogenized using a bead mill (Qiagen TissueLyser LT). RNA was purified using the Qiagen RNeasy Mini kit with DNase (Macherey-Nagel, #740955.250) treatment. Samples were shipped on dry ice to Novogene Europe (UK), where RNA sequencing and bioinformatic analyses were performed.

### Phosphoproteomics

*Drosophila:* Cholesterol *vs.* protein refeeding: Refeeding experiments were conducted as described above in the *Drosophila* refeeding method section. Fifteen (cholesterol assays) or twenty (protein assays) feeding late-third-instar larvae for each of 2 replicates were collected by washing with water, blotted dry on a KimWipe, and snap-frozen in Eppendorf tubes on dry ice. A sample was collected from the nutrient-deprivation condition at the start of refeeding as time-point 0, and further samples were collected at 15 and 30 minutes and at 1, 2, 4, 6, and 10 hours after the start of refeeding. Samples were stored at −80 °C until they were processed. *HR3* and *TOR* knock-down: *Tor* and *HR3* were knocked down in *Drosophila* larvae using the temperature-inducible ubiquitous driver *Tub-GAL80^TS^; Tub-GAL4*. After an 8-hour egg collection at 18 °C, the plates were maintained at 18 °C for an additional 40 hours before first-instar larvae were transferred to vials of STD-FF. These were maintained at 18 °C for an additional 72 hours before being transferred to 29 °C to induce RNAi at 120 h AEL. Nine hours later, at 129 h AEL, the animals were transferred to low-cholesterol synthetic medium (1.2 µg/mL CH) and incubated for 11 further hours at 29 °C before being transferred (at 150 h AEL) to fresh synthetic food containing either 1.2 (continued low-CH exposure) or 80 (replenishment) µg/mL cholesterol, still at 29 °C. After one hour and six hours, two samples of 15 larvae each for each treatment were collected, washed with DI water, blotted dry, and snap-frozen on dry ice before storage at −80 °C.

Karpas-707H: Cells treated for knockdown of *RORα* or *Luciferase* (mock knockdown control) were grown in 6-well plates till 80-90% confluence. Cells were cholesterol-depleted with MCD for 2 hours and washed twice with RPMI. Cholesterol-depletion or cholesterol-stimulation medium was added, and both treatments were sampled in triplicate after one hour; an additional triplicate sample of cholesterol-stimulated cells was taken at 6 hours post-stimulation. All samples were washed once in RPMI. Cells in suspension were collected in a Falcon tube, and cells attached to the culture ware were gently collected in RPMI medium using a cell scraper and added to the suspended cells. Cells were pelleted at 900 RCF for 4 minutes, supernatant was removed, and cells were frozen on dry ice and stored at −80 °C.

Proteins from larval samples were extracted in 300 µL 5% sodium deoxycholate (SDC) in 50 mM HEPES (pH 8.5) containing Roche cOmplete protease inhibitor and PhosSTOP phosphatase inhibitors (Sigma) using a FastPrep-24 bead beater (MP Biomedicals) using 25-30 1.4 mm ceramic beads (OMNI international, US). Each tube was subjected to 3×45-second bead beating. After treatment the solution was diluted with an additional 300 µL of 50 mM HEPES (pH 8.5) and subjected to a second round of 3×45-second bead beating. The samples were centrifuged, and the supernatant was transferred to a low-binding 1.5-mL Eppendorf tube and subjected to probe sonication for 2×20 seconds at 60% amplitude. After sonication, the sample was denatured at 110 degrees for 5 minutes and centrifuged for 20 min at 20,000x*g* to pellet insoluble material. The supernatant was transferred to another tube and the protein concentration was measured using a Nanodrop spectrophotometer. For the Karpas 707h cells, the proteins were extracted from the cell pellets in 300 µL 3% SDC in 50 mM HEPES, pH 8.5, containing cOmplete protease inhibitor and PhosSTOP phosphatase inhibitors (Sigma), using probe sonication for 2×20 seconds at 60% amplitude. After sonication, the sample was denatured at 110 degrees for 5 minutes and subsequently centrifuged for 20 minutes at 20,000x*g* to pellet insoluble material. The supernatant was transferred to another tube and the protein concentration was measured using a Nanodrop N60 spectrophotometer.

A total of 100 µg of protein was taken from each sample and subjected to reduction and alkylation using 10 mM DTT for 20 minutes followed by 20 mM iodoacetamide for 20 minutes. Trypsin (5%) was added and the solutions were incubated at 37 °C overnight. After incubation, a further 1% bolus of trypsin was added and the samples were incubated for one further hour at 37 °C. After incubation, the cleaved peptide solutions were labeled with tandem mass tags (TMTpro) 18-plex according to the manufacturer’s protocols. After labeling, the 18 samples were combined into one sample containing all the larval peptides and one containing the Karpas 707h material. The SDC was removed by acidification and subsequent centrifugation for 20 minutes at 20,000x*g*. The supernatant was transferred to a low-binding Eppendorf tube and dried until 150 µL was left. The enrichment of phosphopeptides and non-modified peptides using titanium dioxide and subsequent high-pH reversed-phase (RP) fractionation were performed as described^71^.

The phosphopeptide and non-modified-peptide fractions from the larvae and Karpas 707h cells were analyzed by tandem mass spectrometry using an EASY nanoLC system coupled with a Fusion Lumos Tribrid or an Orbitrap Eclipse Tribrid. Lyophilized peptides from the high-pH RP fractionation (12-20 concatenated fractions) were re-solubilized in 3-5 µL of 0.1% formic acid (FA) and loaded onto a 20-cm analytical column (100-μm inner diameter) packed with ReproSil – Pur C18 AQ 1.9 μm RP material. The peptides were eluted with an organic solvent gradient from 100% phase A (0.1% FA) to 25% phase B (95% ACN, 0.1% FA) for 80-100 minutes (depending on fraction), then from 25% B to 40% B for 10-20 minutes before the column was washed with 95% B. The flow rate was set to 300 nL/minute during elution. For the two instruments the automatic gain-control target value of 1.5×10^6^ ions in MS and a maximum fill time of 50 ms were used. Each MS scan was acquired at high resolution (120,000 full width half maximum (FWHM)) at *m/z* 200 in the Orbitrap with a mass range of 350-1400/1500 Da. The instruments were set to select as many precursor ions as possible in 3 seconds between the MS analyses. Peptide ions were selected from the MS for higher-energy collision-induced dissociation (HCD) fragmentation (collision energy: 34%). Fragment ions were detected in the Orbitrap at high resolution (50,000 FWHM) for a target value of 1.5×10^5^ ions and a maximum injection time of 200 ms (phosphopeptides) or 86 ms (“non-modified peptides”) using an isolation window of 0.7 Da and a dynamic exclusion of 20-45 seconds. All raw data were viewed in Xcalibur v3.0 (ThermoFisher Scientific).

### Peptide/protein identification and quantitation

All LC-MS/MS raw data files from *Drosophila*-larva experiments were searched in Proteome Discoverer (PD) version 2.5.0.400 (ThermoFisher Scientific). The raw data were searched in PD using the SEQUEST HT search algorithm against the Fly Database protein FASTA file. The searches had the following criteria: enzyme, trypsin; maximum missed cleavages, 2; fixed modifications, TMTpro (N-terminal), TMTpro (K) and Carbamidomethyl (C).

Variable modification for the phosphopeptides was Phospho (S/T/Y) and Deamidation (N) whereas no variable modifications were used for the “non-modified peptides”. For the results from the Karpas 707h cells the raw data files were searched in PD using first an in-house Mascot server (Version 2.2.04, Matrix Science Ltd., London, UK) against the Swissprot human protein database using the following criteria: enzyme, trypsin; maximum missed cleavages, 2; fixed modifications, TMTpro (N-terminal), TMTpro (K) and Carbamidomethyl (C). Variable modification for the phosphopeptides was Phospho (S/T/Y) whereas no variable modifications were used for the “non-modified peptides”. The peptide fragment ion spectra that were not identified with a peptide in Mascot with below 1% False Discovery Rate (FDR) was further subjected to database searching using SEQUEST HT in PD against the Human Uniprot FASTA database, using the same criteria as above. The TMTpro reporter ion signals were quantified using S/N and they were normalized to the total peptide S/N in the PD program. The in-built ANOVA test in PD was used to generate *p*-values for all the phosphopeptides and proteins identified in the database searches.

### Data filtering and bioinformatics analysis of phosphoproteomics and RNAseq

Phosphoproteomics data for pathway analysis was filtered for differences >30% in magnitude and *p*<0.05 between cholesterol starvation and replenishment within each genotype. An Excel macro was used to identify populations of interest. For the RNAseq experiments, gene-expression levels are reported as FPKM (fragments per kilobase of transcript per million bases sequenced), which takes into account both sequencing depth and gene length. Differential expression analysis was conducted by Novogene by read-count normalization, model-dependent *p*-value estimation, and false-discovery-rate estimation based on multiple hypothesis testing. Differential expression was further analyzed by comparing genotype-based differences between sets of genes exhibiting cholesterol-induced expression change, resulting in 7 categories of genotype-specific cholesterol-induced regulation (with set sizes from Figure 5D for illustration): (1) genes requiring both HR3 and TOR to exhibit a cholesterol-induced change (occurring only in the control genotype) – 980; (2) those cholesterol-induced changes that require HR3 but are independent of Tor (occurring in controls and *Tor* knockdowns) – 452; (3) cholesterol-induced changes requiring TOR but independent of HR3 (occurring in controls and *HR3* knockdowns) – 374; (4) those changes requiring neither HR3 nor TOR (occurring in all samples) – 267; (5) cholesterol-induced changes repressed by HR3 but not by TOR – novel changes occurring only in *HR3-RNAi* samples – 1029; (6) cholesterol-induced changes repressed by TOR but not by HR3 – novel novel changes occurring only in *TOR-RNAi* – 644; (7) cholesterol-induced changes repressible by either HR3 or TOR – novel changes that occur in both knockdowns but not in controls – 125. Pathway analysis was carried out on both the phosphoproteomics and RNAseq data using the Panther Classification system (Protein Analysis Through Evolutionary Relationships), with the statistical overrepresentation test using Reactome database (version 85)^68^. In addition to the statistical analysis already conducted during data filtering, a Fisher’s t-test was applied, and only pathways with a *p*<0.05 for enrichment were considered. For RNA-seq data, model-dependent *p*-values were used, and for all phosphopeptides, *p*-values were determined using the ANOVA test in Proteome Discoverer.

### Recombinant HR3 protein expression and purification

For baculovirus-mediated expression in insect cell culture, the HR3 LBD (I230-T487; GenBank accession NP_788303) was subcloned into a modified pFASTBAC DUAL vector (Invitrogen) that contained a 6xHis tag and a thrombin cleavage site as described^72^. The hexahistidine-tagged protein was expressed in Hi5 insect cells (*Trichoplusia ni*) grown in 1 liter of M3 medium (Sigma) supplemented with 10% fetal bovine serum (Wisent). Cells were grown at 26 °C for two days in baffled flasks in a shaking incubator. The cells were harvested by centrifugation, washed with PBS (20 mL), and centrifuged again before the pellets were frozen in liquid nitrogen and stored at −80 °C. Prior to purification, the cell paste was thawed, resuspended in lysis buffer, and sonicated on ice 3 times for 1 minute each. Lysates were clarified by centrifugation, and tagged protein was bound using Ni-NTA affinity chromatography (column volume 1 mL). Column-bound protein was washed with 300 mL of wash buffer (30 mM imidazole, 500 mM NaCl, 5% glycerol, 0.5 mM TCEP, and 10 mM Tris; pH 8.2). Protein was eluted from the column using a similar buffer containing 250 mM imidazole. The eluate was concentrated to a volume of 5 mL and loaded onto a size-exclusion-chromatography column (Superdex 16/60, GE Healthcare); HR3 LBD was eluted in a buffer of 150 mM NaCl, 0.5 mM TCEP, and 10 mM Tris (pH 8.2) using an AKTA FPLC apparatus (GE Healthcare). After using SDS-PAGE to confirm the purity of eluted fractions, a 10-mg sample of the receptor protein was diluted in buffer Q_ο_ (30 mM NaCl, 5% propanediol, 5 mM DTT, 20 mM Tris, pH 8.5) and loaded onto a Source 30Q anion-exchange column (GE Healthcare). Using a linear NaCl gradient (30-mM to 2-M NaCl in Q_ο_), the protein was eluted using an AKTA FPLC (GE healthcare) and again checked for purity using SDS-PAGE.

### Non-denaturing mass spectrometry

ES-MS was carried out as previously described^72^. Briefly, mass analysis was done using a quadrupole time-of-flight mass spectrometer (Q-TOF, Micromass) equipped with a nano electrospray (ES) source. Purified HR3 LBD was desalted in four dilution/concentration steps using centrifugal concentration (Millipore) into 20-mM ammonium acetate (pH 6.2) and adjusted to a protein concentration of 1 mg/ml. Protein was loaded in a gold-coated capillary (Protona) with the tip opened to produce an orifice of approximately 10 μm. Positive ESI-MS was performed at a capillary voltage of 1.8 to 2 kV and a cone voltage of 10–50 V. Collision-induced dissociation experiments were carried out with a collision energy of 80–150 V using argon as the collision gas. Average molecular masses were calculated using Maxnet1 in Mass Lynx 4.0 (Micromass).

### Extraction of HR3 and GC/MS analysis

Using solvent-washed glassware, endogenous ligand was extracted from 1 mg of purified HR3 LBD by combining 3 mL of purified receptor with 9 mL of acidified chloroform:methanol (2:1 v/v) and mixing vigorously for 1 minute. Organic and aqueous phases were separated by centrifugation, and the organic phase was transferred to a clean tube. The aqueous phase was extracted twice more, and the organic fractions were pooled. The acidified chloroform:methanol extract was repeatedly washed with 0.2 volumes of distilled water and reseparated until the aqueous phase became neutral (pH 7). The organic extract was evaporated to dryness under nitrogen, and residue was dissolved in 50 μL pyridine. A 20-μL aliquot of this sample was combined with 10 μL of N,O-bis-(trimethylsilyl)trifluroacetamide (N,O-BSTFA) and trimethylsilyl-derivatized at 60 °C for 30 minutes. The same procedure was followed for derivatization of the cholesterol reference^73^ (Steraloids).

### Gas chromatography mass spectrometry

The derivatized samples were analyzed using a gas chromatograph (3900 GC, Varian) coupled to an ion-trap mass spectrometer (Saturn 2100T MS/MS, Varian) equipped with an electron-impact ion source. Samples were chromatogaphically separated using helium as a carrier gas (1.5 mL/min) on a 30-m x 0.25-mm x 0.25-μm fused-silica column (CP8944, Varian) using a two-segment temperature gradient of 80-150 °C at 20 °C/minute and 150-345 °C at 10 °C/minute. Eluted analyte from chromatography was ionized in the positive-ion mode with a scan range of 40–650 *m/z*. Data was analyzed using MS Data Review in MS workstation (Varian).

### HR3 Ligand-sensor embryo treatment

Transgenic HR3 ligand-sensor (*HR3::GAL4; UAS-GFP*) embryos were permeabilized and treated for 15 minutes at 25 °C with cholesterol in MBIM culture media as previously described in Palanker *et al.*^34^ The MBIM was then removed, and the embryos covered with halocarbon oil and allowed to develop for a minimum of 2 hours prior to observation.

### Statistical analysis

Statistical analysis was computed using the Prism software package (GraphPad, version 10). Fat-body pS6 data was tested for outliers using the ROUT method and a Q of 10 due to the large variation within the tissue. All data sets were assessed for normality prior to statistical analysis, and appropriate parametric or non-parametric tests were selected. Graphs were generally created using the Prism software, showing all data points and mean ± standard error of the mean. Detailed statistical information can be found in each figure’s legend. Graphical illustrations of RNAseq and phosphoproteomics data were made using SRplot^69^.

## Acknowledgements

This work was supported by funding from the Danish Independent Research Council – Natural Sciences (8021-00055B) to KR. TK and KVH were supported by funding from the Danish Independent Research Council – Natural Sciences (9064-00009B) to KVH. The Zeiss LSM 900 confocal microscope and the PerkinElmer Ensight plate reader were supported by infrastructure grants from the Carlsberg Foundation (CF19-0353 and CF17-0615) to KR et al.

## Author contributions

ML, MJT, KP, HK, and KR conceived and designed the study. ML, KP, LHP, OK, TK, SN, SL, AK, GL, AE, AAT, MRL, HMK, MJT, and KR designed, performed, and analyzed experiments. ML, MJT, and KR wrote the manuscript.

## Competing interests

Authors declare that no competing interests exist.

## Supplemental Information

**Figure S1.**
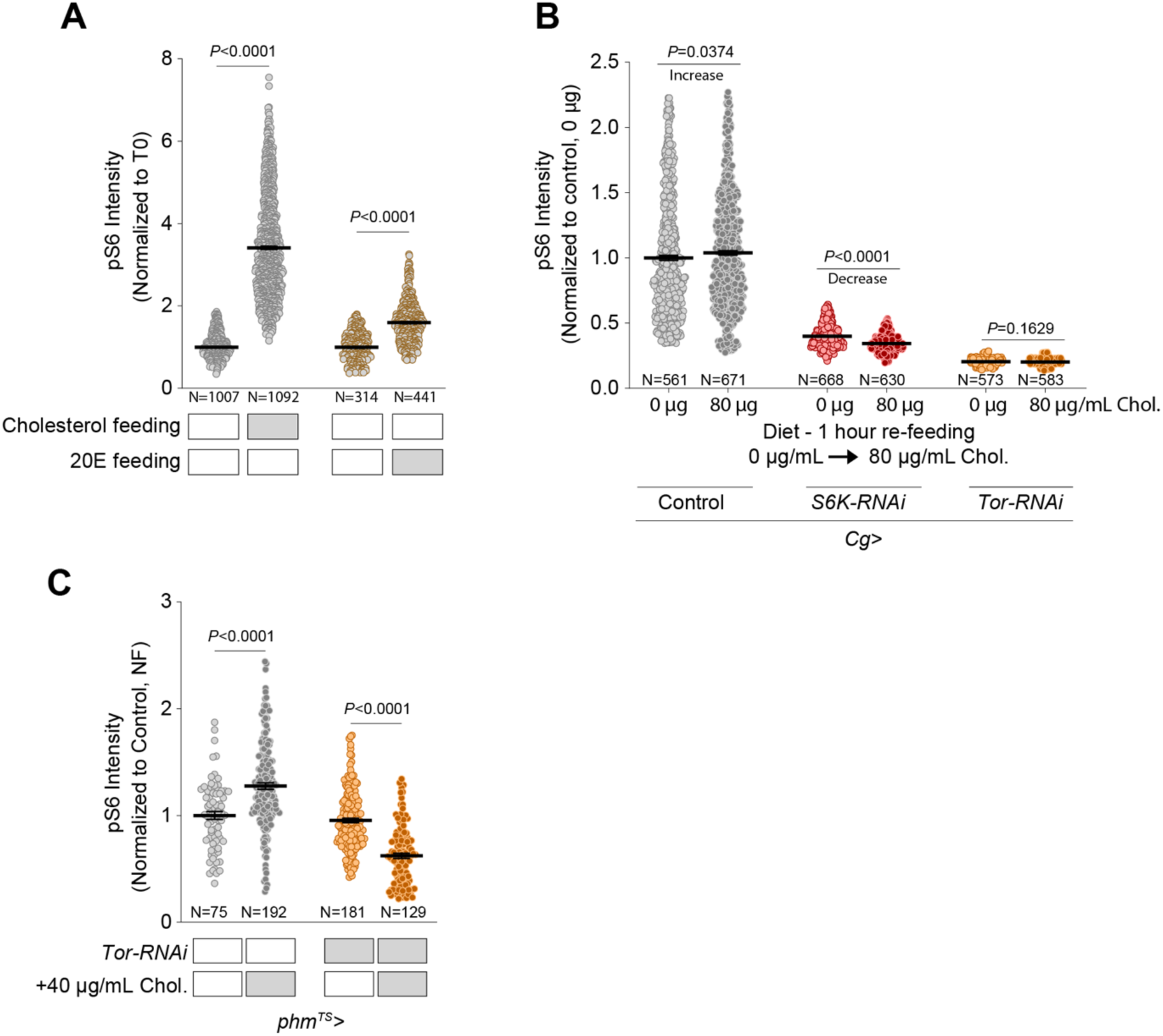
(A) Quantification of wild-type larval fat-body pS6 response to 6 hours’ feeding with cholesterol (80 µg/mL) or 20-hydroxyecdysone (250 µg/mL with 1.2-µg/mL cholesterol) following 10-hour low-cholesterol (1.2 µg/mL) feeding. (B) Phospho-S6 staining of fat-body tissue from controls and animals expressing fat-body-specific knock-down of *S6K* or *Tor*, after 1 hour’s refeeding with 0- or 80-µg/mL-cholesterol medium following 10-hour cholesterol deprivation (0 µg/mL cholesterol in synthetic medium). (C) Phospho-S6 response in PG cells of control larvae and animals expressing PG-specific *Tor* knockdown, chronically fed NutriFly or NutriFly supplemented with 40 µg/mL cholesterol. Statistics: (A, B, C) All data plotted as means ±SEM, normalized to control cholesterol-starvation condition (T0). Each data point represents pS6 intensity in the cytoplasm of a single cell. *P*-values calculated by Kruskal-Wallis ANOVA tests with Dunn’s multiple comparisons.

**Figure S2.**
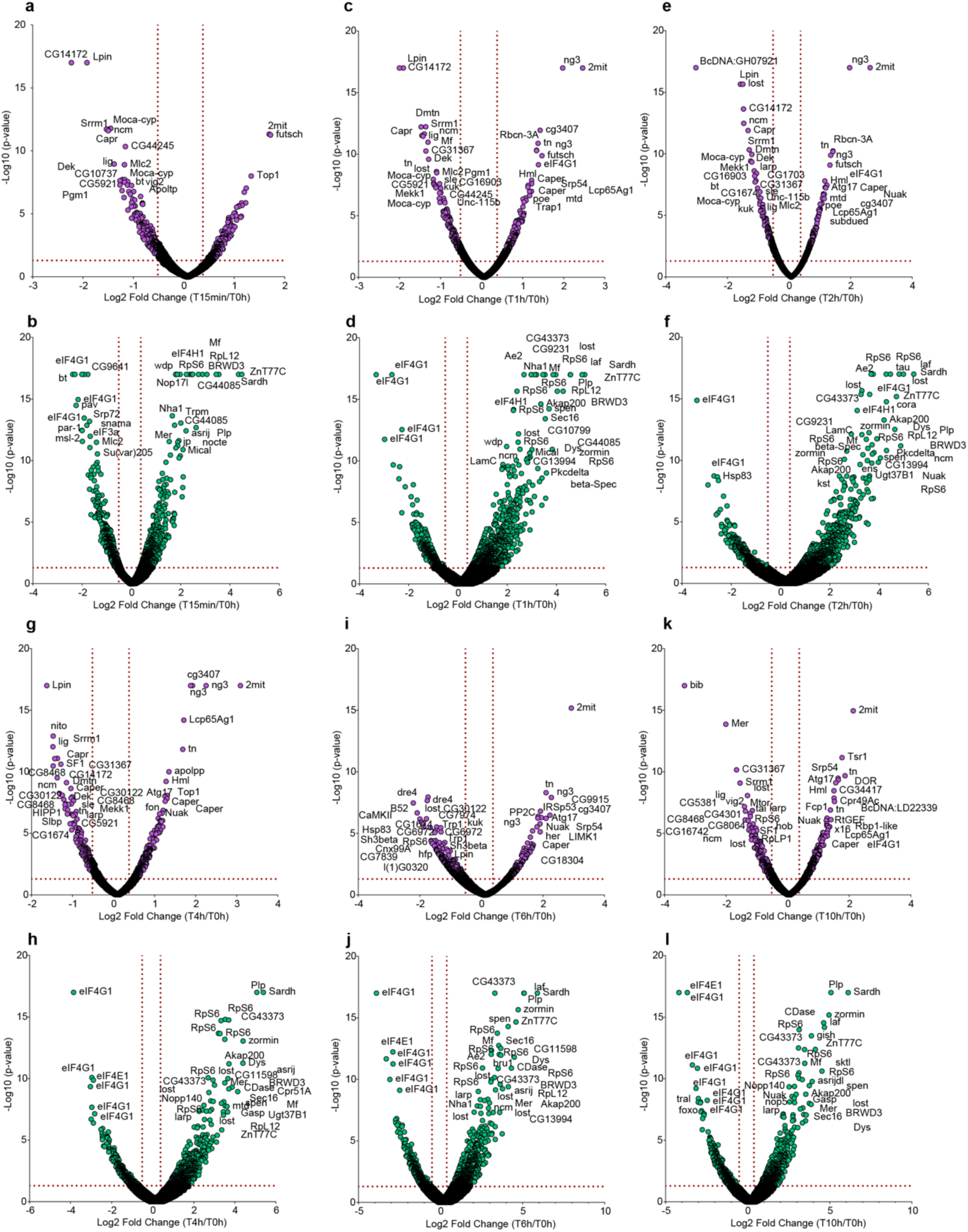
(A, C, E, G, I, K) Volcano plots of phosphoproteomic changes in response to cholesterol refeeding (80 µg/mL) following 10-hour cholesterol deprivation (0 µg/mL), in animals harvested after 15 minutes or 1, 2, 4, 6, or 10 hours. (B, D, F, H, J, L) Similar plots for animals refed with protein (14.7 mg/mL) following 10-hour protein deprivation. Statistics: Differences in phosphorylation state were considered significant if they were larger than 30% (log_2_ fold change x>0.379 or x<-0.515) with *p*<0.05 (-log_10_>1.3), thresholds marked with dotted red lines.

**Figure S3.**
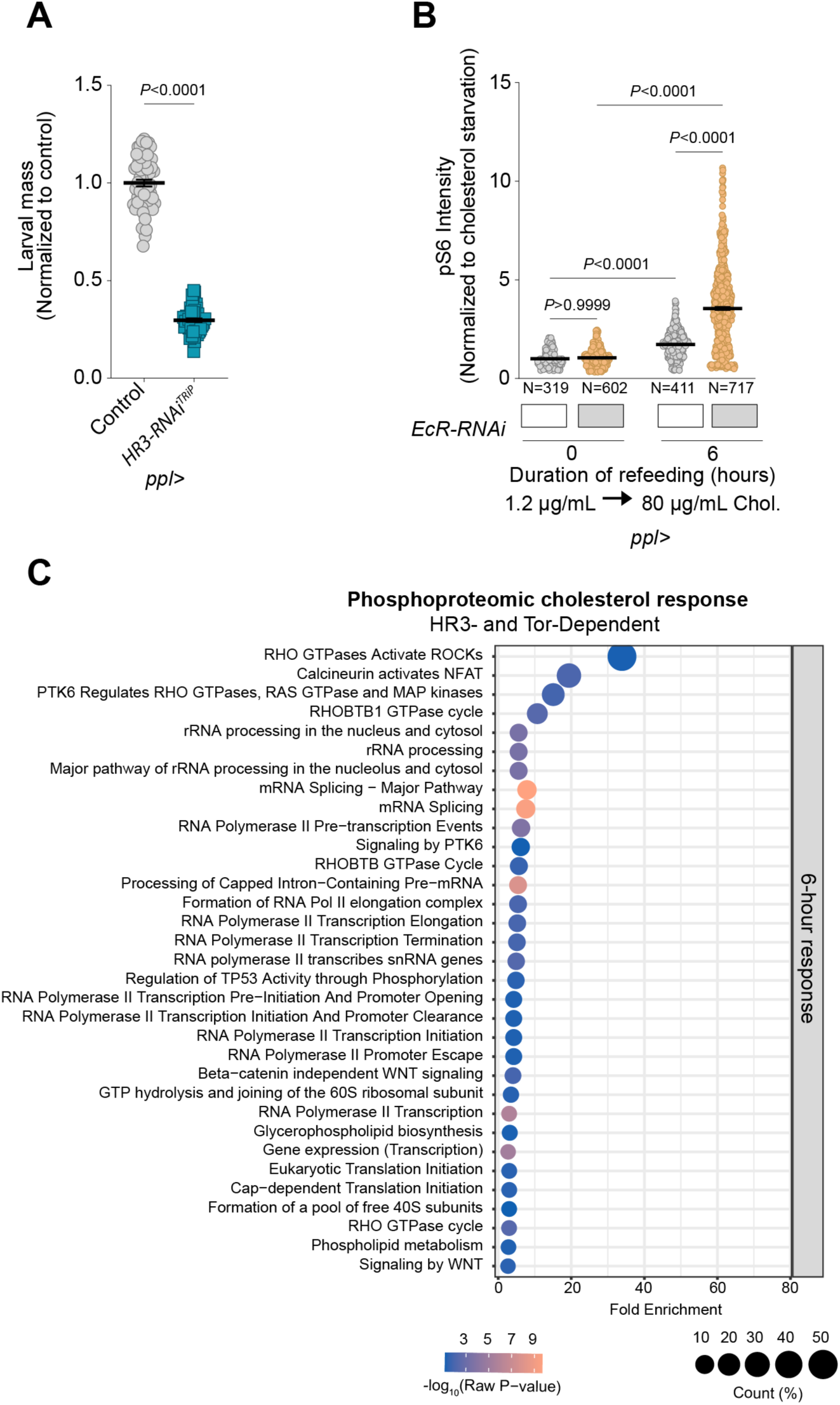
(A) Larval mass at 96 hours AEL of controls and animals expressing an additional *HR3* knock-down construct in the fat body using *ppl>*, fed on STD-FF. (B) Fat-body pS6 response to cholesterol refeeding (80 µg/mL cholesterol) in controls and larvae expressing *EcR* knock-down in the fat-body with *ppl>*, following 10-hour feeding with low-cholesterol (1.2 µg/mL) medium. Each data point reflects the cytoplasmic pS6 intensity of a single cell (C) Selected pathways enriched in proteins exhibiting phosphoproteomic responses to 6-hour cholesterol refeeding that were dependent on both HR3 and Tor. Pathway analysis was conducted using Panther and Reactome. Statistics: (A, B) Data plotted as means ± SEM. (A) *P*-value calculated using two-tailed unpaired t-test. (B) *P*-values calculated using Kruskal-Wallis ANOVA test with Dunn’s multiple comparisons. (C) Only pathways with *p*<0.05 by Fisher’s t-test are included.

**Figure S4.**
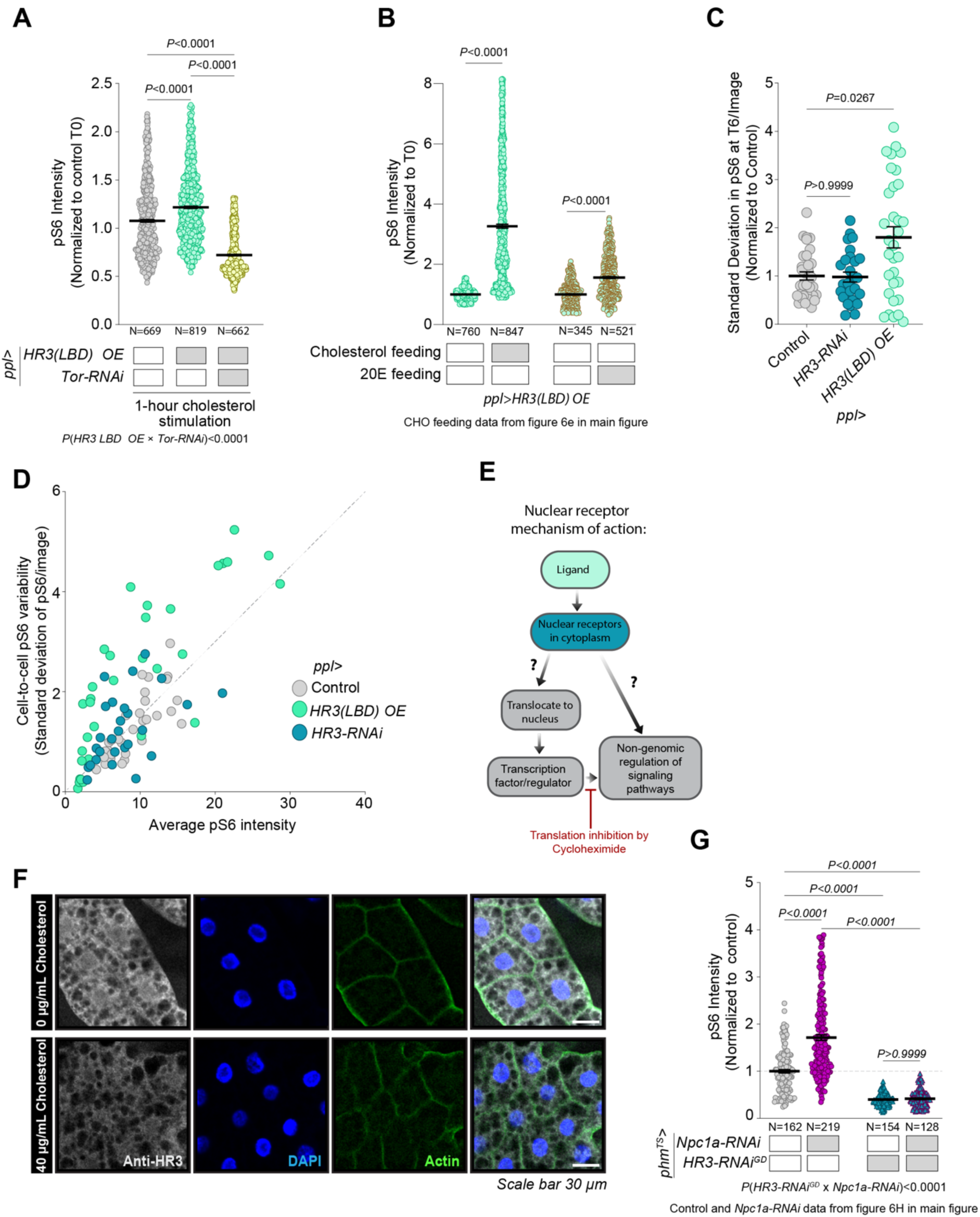
(A) Fat-body pS6 response to 1-hour cholesterol refeeding (80 µg/mL) in controls and animals overexpressing HR3(LBD) in the fat body using *ppl>*, with and without simultaneous *Tor* knockdown, following 10-hour feeding with low-cholesterol (1.2 µg/mL) medium. (B) Phospho-S6 responses in fat-body cells of larvae overexpressing HR3(LBD) in the fat body (*ppl>*) to feeding with cholesterol (80 µg/mL) or 20-hydroxyecdysone (20E, 250 µg/mL with 1.2 µg/mL cholesterol) for 6 hours following 10-hour feeding on low-cholesterol (1.2 µg/mL) medium. (C) Quantification of cell-to-cell variation in pS6 intensity. (D) Cell-to-cell variability in pS6 signal in fat-body tissue fragments of control larvae and animals either expressing *HR3* knockdown or overexpressing HR3(LBD) in the fat-body (*ppl>*) after 6 hours’ cholesterol re-feeding (80 µg/mL) following 10-hour low-cholesterol feeding (1.2 µg/mL). (E) Illustration of two potential mechanisms of TOR regulation by nuclear receptors such as HR3, illustrating the impetus for the *ex-vivo* experiments. Treatment with cycloheximide inhibits one of the two routes by inhibiting translation; thus the fraction of cholesterol-induced pS6 increase remaining with cycloheximide treatment is due to non-genomic effects on the Tor signaling pathway. (F) Representative immunostains against HR3 in larval fat-body tissue after cholesterol starvation (6 hours) and after cholesterol re-feeding (0 µg/mL for 10 hours). Scale bar is 30 µm. (G) pS6 staining in the prothoracic gland with *Npc1a-RNAi* alone, and in combination with *HR3* knockdown. Statistics: Data are plotted in (A, B, C, G) as means with SEM. *P*-values calculated using Kruskal-Wallis test with Dunn’s multiple comparisons and two-way ANOVA test for interactions. Data are normalized to *phm^TS^>* driver control, graphed as mean ± SEM. (A) Data are normalized to control. Kruskal-Wallis with Dunn’s multiple comparisons and a two-way ANOVA test for epistasis. (B) Data are normalized to dietary controls (T0). Kruskal-Wallis ANOVA with Dunn’s multiple comparisons. (C) Each point represents the brightness and variation within one piece of fat-body tissue, such as those in panel. (D) Each data point reflects the standard deviation among all the single-cell pS6 measurements in a single analyzed tissue fragment.

## Notes

### Competing Interest Statement

The authors have declared no competing interest.

